# Pan-cancer circular genomics identifies intratumoral *Staphylococcus lugdunensis* as a metabolic driver in bladder cancer

**DOI:** 10.64898/2026.04.14.717102

**Authors:** Tao Wang, Ming Zhao, Conghui Li, Zhuofan Bai, Fan Zhang, Hongyu Zhao, Wei Lv, Chunhua Lin, Duanyang Kong, Xiaolu Zhao, Pengcheng Wang, Fengbiao Mao

**Affiliations:** Institute of Medical Innovation and Research, Peking University Third Hospital, Beijing, China; Cancer Center, Peking University Third Hospital, Beijing, China; Beijing Key Laboratory for Interdisciplinary Research in Gastrointestinal Oncology (BLGO), Beijing, China; Department of Medicine Huddinge, Center for Hematology and Regenerative Medicine, Karolinska Institutet, Stockholm, Sweden; State Key Laboratory of Chemical Resources Engineering, Beijing University of Chemical Technology, Beijing, China; Department of Urology, Peking University Third Hospital, Beijing, 100191, China; State Key Laboratory of Female Fertility Promotion, Center for Reproductive Medicine, Department of Obstetrics and Gynecology, Peking University Third Hospital, Beijing, China; National Clinical Research Center for Obstetrics and Gynecology (Peking University Third Hospital), Beijing, China; Key Laboratory of Assisted Reproduction (Peking University), Ministry of Education, Beijing, China; Beijing Key Laboratory of Collaborative Innovation in Frontier Technologies for Population Quality, Beijing, China; Hangzhou Institute of Medicine (HIM), Chinese Academy of Sciences, Hangzhou, Zhejiang, China; Division of Immunology, Department of Medical Biochemistry and Biophysics, Karolinska Institutet, Stockholm, Sweden; The Affiliated Yantai Yuhuangding Hospital of Qingdao University, Yantai, Shandong, China

**Keywords:** Intratumoral bacteria, Bladder cancer, Circular genomics, *Staphylococcus lugdunensis*, LPC14:0, PPARδ, Lipid metabolism

## Abstract

Intratumoral microbiota play key roles in cancer but are challenging to profile due to low biomass. We developed CIRCMIP, an enzymatic pipeline that eliminates linear DNA to enrich bacterial genomes, achieving 100-fold higher sensitivity than standard metagenomics. Applied to 312 pan-cancer specimens, CIRCMIP identified *Staphylococcus lugdunensis* as a signature bacterium enriched in early-stage bladder cancer (BLCA), where its presence predicts poor survival. Integrated modeling and lipidomics revealed that *Staphylococcus lugdunensis* colonization drives aberrant lipid metabolism with secretion of LPC14:0. Both *Staphylococcus lugdunensis* and LPC14:0 drive BLCA progression by promoting fatty acid uptake and β-oxidation. Mechanistically, chemical proteomics revealed LPC14:0 as a direct PPARδ ligand, binding via hydrogen bonds with Thr292/Thr289. This activation upregulates fatty acid transporters (*CD36*, *FABP4*) and metabolic enzymes (*ACOX2*), fueling malignant proliferation. Furthermore, CIRCMIP-derived biomarkers show robust diagnostic accuracy, establishing a new research paradigm and revealing the *Staphylococcus lugdunensis*-LPC14:0-PPARδ axis as a therapeutic target in bladder cancer.

**HIGHLIGHT:**

- CIRCMIP leverages DNA circularity to achieve 100-fold sensitivity in pan-cancer bacteriome profiling.
- *Staphylococcus lugdunensis* is a bacterial signature in early-stage bladder cancer that predicts poor patient survival.
- Bacteria-derived LPC14:0 acts as a direct ligand for PPARδ to drive host malignant metabolic reprogramming.
- Targeting the *Staphylococcus lugdunensis*-LPC14:0-PPARδ axis abrogates malignant progression in bladder cancer.

## INTRODUCTION

Cancer represents a formidable global health crisis, accounting for millions of deaths annually. Notably, over 16% of cancer cases are etiologically linked to infectious agents, highlighting the critical interplay between microbial communities and oncogenesis^1^. While the gut microbiome has been extensively studied for its role in modulating tumor progression, metastasis, and therapeutic responses^2–9^, emerging evidence has unveiled intratumoral bacteria as a previously underappreciated component within the tumor microenvironment (TME)^10^, a complex ecosystem where diverse cellular components and molecular drivers orchestrate malignancy^11–14^. These microbial inhabitants are increasingly implicated in core cancer hallmarks, including immune evasion, metabolic reprogramming, and therapeutic resistance^15^. Characterizing their functional contributions not only advances our understanding of tumor biology but also opens new avenues for precision diagnostics, prognostics, and microbiome-targeted therapies^16,17^.

The intratumoral microbiome exhibits remarkable specificity, with distinct taxonomic profiles demarcating tumor types and subtypes. For instance, Proteobacteria (Pseudomonadota) dominate pancreatic ductal adenocarcinoma, whereas colorectal carcinomas are enriched with Firmicutes (Bacillota) and Proteobacteria^15^. These tumor type-associated microbial signatures hold promise as diagnostic biomarkers, particularly for malignancies lacking reliable molecular markers. Furthermore, certain bacteria exhibit selective tropism for tumor tissues, a property exploitable for targeted drug delivery. Perhaps most compellingly, longitudinal studies reveal correlations between microbial community composition and patient survival, positioning the intratumoral microbiome as a prognostic tool^15^. However, translating these insights into clinical practice is impeded by persistent methodological and biological challenges.

A central obstacle lies in the extremely low biomass of intratumoral bacteria, which often constitute <0.01% of total DNA in tumor samples. This scarcity exacerbates contamination risks from environmental sources or DNA extraction kits^18–20^, particularly in large-scale collaborative efforts like The Cancer Genome Atlas (TCGA). Although TCGA-based analyses linked pathogens such as *Fusobacterium nucleatum* (*F. nucleatum*) to colorectal cancer progression^3,21–24^, distinguishing endogenous microbial signals from artifacts remains contentious^25^. While 16S rRNA sequencing provides cost-effective taxonomic profiling, its resolution is limited to genus-level classification and prone to PCR amplification biases. Shotgun metagenomics enables strain-level characterization but requires high sequencing depth to overcome host DNA interference (>99% of reads in clinical samples). Emerging technologies, including single-cell sequencing and spatial transcriptomics, offer partial solutions by isolating bacterial cells or mapping their spatial niches within tumors. Yet, these methods remain technically demanding and lack scalability for large cohorts. Mechanistically, intratumoral bacteria engage in multifaceted crosstalk with host cells. For example, *F. nucleatum* in colorectal cancer activates autophagy pathways to confer chemoresistance, while *Bifidobacterium* enhances anti-PD-1 immunotherapy efficacy by promoting dendritic cell activation^3,22^. Recent studies have further demonstrated that the specific spatial engagement between bacteria and cancer cells dictates divergent tumor immune responses^26^. Furthermore, tumor-resident *Staphylococcus* has been shown to drive metastatic colonization in lung adenocarcinoma by secreting lactate to remodel the metabolic landscape of the tumor niche^27^. These findings underscore the urgency of deciphering host-microbe interaction networks through multi-omics integration. However, existing methodologies lack the sensitivity to resolve rare taxa or distinguish functionally active bacteria from DNA remnants, leaving the metabolic influence of the intratumoral microbiome largely overlooked. Specifically, while host-intrinsic factors have been implicated in stimulating fatty acid metabolism to promote bladder cancer (BLCA) progression^28^, the role of tumor-resident microbiota as an exogenous driver of this metabolic shift remains largely unexplored.

In this study, we utilize CIRCMIP to systematically map the pan-cancer intratumoral bacteriome by selectively capturing circular bacterial genomes. This high-resolution profiling identifies *Staphylococcus lugdunensis* (*S. lugdunensis*) as a signature symbiont specifically enriched in bladder cancer (BLCA), where its abundance serves as a potent predictor of poor prognosis. We demonstrate that *S. lugdunensis* drives malignant progression by secreting LPC14:0, which acts as a direct ligand for host PPARδ. This interaction triggers a transcriptional program that heightens fatty acid uptake and β-oxidation, thereby fueling tumor cell proliferation. Our findings establish the *S. lugdunensis*-LPC14:0-PPARδ axis as a decisive factor in BLCA metabolic reprogramming and highlight its potential as a targetable microbial-metabolic signaling pathway. Moreover, by overcoming the fundamental constraints of existing methodologies, CIRCMIP establishes a new paradigm for high-fidelity intratumoral bacteriome research, providing a scalable platform for systemic biomarker discovery. Ultimately, our study not only elucidate the functional roles of tumor-resident bacteria in metabolic oncogenesis but also offers novel avenues for therapeutic development.

## RESULTS

### CIRCMIP empowers high-confidence characterization of the tumor bacteriome

To overcome the inherent challenges of profiling intratumoral bacteria, which are frequently obscured by overwhelming host DNA backgrounds, we exploited the distinctive circular topology of bacterial genomes to develop CIRCMIP (CIRCular DNA-based Microbiome Identification Pipeline). This methodological framework employs an enzymatic enrichment strategy to selectively capture circular bacterial DNA, effectively purging host genomic interference and overcoming the low-biomass constraints of the tumor microenvironment (Figure 1A). CIRCMIP integrates a multi-stage analytical workflow encompassing stringent host DNA filtration, high-confidence species identification, and machine learning-driven biomarker discovery (see Methods). We applied CIRCMIP to a pan-cancer cohort of 312 samples across eight distinct malignancies, including bladder cancer (BLCA), colorectal cancer (COADREAD), liver hepatocellular carcinoma (LIHC), lung adenocarcinoma (LUAD), stomach adenocarcinoma (STAD), prostate adenocarcinoma (PRAD), ovarian carcinoma (OV), and medulloblastoma (MB) (Table S1). This approach successfully recovered an abundance of bacterial genomes and plasmids derived from circular DNA, whereas linear DNA-associated fungal sequences remained virtually undetectable (Figures 1B, S1A, and S1B; Table S2). Notably, CIRCMIP exhibited markedly enhanced sensitivity, achieving a more than 100-fold increase in bacterial read detection compared to conventional whole genome shotgun (WGS) metagenomics. This approach consistently yielded superior performance in both bacterial biomass quantification and the characterization of taxonomic richness across the samples (*P* < 0.01) (Figure 1C; Table S2). To ensure the highest degree of taxonomic accuracy, we implemented a rigorous five-tiered filtration pipeline to refine our initial census of 7,283 putative bacterial species. This stringent funneling process, which integrated library contaminant removal, sample-level noise reduction, benchmarking against 2,491 sterile cell-line samples, normal-tissue background subtraction, and low-abundance taxon filtering, distilled the dataset into 732 high-confidence species (Figure S1C; Table S2). Taken together, these results establish CIRCMIP as a robust and highly sensitive framework for the high-resolution characterization of the intratumoral bacteriome.

**Figure 1.**
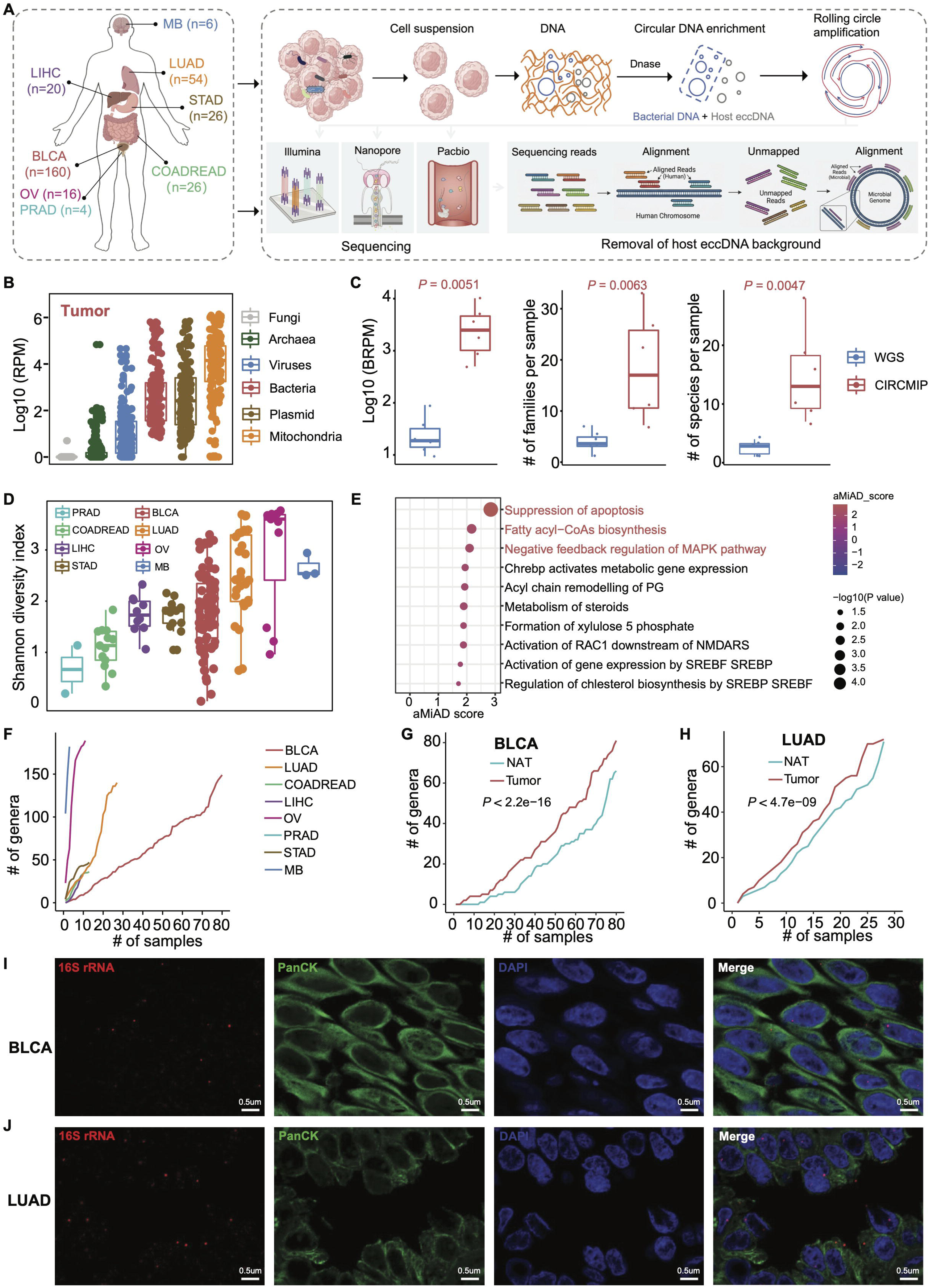
Pan-cancer landscape of the intratumoral bacteriome identified via CIRCMIP. (A) Schematic representation of the CIRCMIP workflow. The pipeline integrated clinical specimen from eight cancer types, utilizing DNase to selectively degrade linear host DNA and facilitate the enrichment of circular genetic templates for rolling circle amplification. Following high throughput sequencing, host circular genetic backgrounds were computationally depleted via alignment to the human reference genome. This integrated approach enabled the high-fidelity identification of bacterial circular DNA signatures within the tumor tissues. (B) Composition of the intratumoral circularome within the pan-cancer cohort. Bacterial (red) and plasmid (brown) DNA represented the predominant genetic components identified across the cohorts, while linear fungal DNA (grey) was minimally detected. RPM, reads per million total reads. (C) Sensitivity benchmarking of CIRCMIP and WGS. The CIRCMIP method achieved significantly higher bacterial read detection and identified a greater number of bacterial taxa compared to conventional whole genome sequencing (*P* < 0.01, wilcoxon test). BRPM, bacterial reads per million total reads. (D) Alpha diversity analysis using the Shannon index to characterize the complexity of bacterial communities across the eight solid tumor types at the species level. (E) Functional association analysis of bacterial signatures with host biological pathways. The X axis represents the aMiAD score derived from Adaptive Microbiome alpha diversity Association Analysis utilizing a Gaussian kernel. A higher aMiAD score indicates a stronger statistical association between the identified bacterial signatures and the respective biological pathways. (F) Genus accumulation curves illustrating the cumulative number of bacterial genera discovered relative to the number of samples analyzed across all cancer types. (G and H) Comparative accumulation analysis between tumor and matched normal adjacent tissues in BLCA (G) and LUAD (H) cohorts (paired t-test). (I and J) Spatial visualization of the intratumoral bacteria via RNAscope combined with immunofluorescence. Representative images from BLCA (I) and LUAD (J) specimens confirmed the presence of bacteria within the tumor tissue. Bacterial 16S rRNA (red) was detected utilizing RNAscope technology, while epithelial tumor cells were identified by PanCK immunostained signals (green). Nuclei were counterstained with DAPI (blue). Scale bars represent 0.5 μm. See also Figure S1 and Table S1, S2.

Leveraging the enhanced sensitivity and specificity of the CIRCMIP framework, we extended our analysis to a broad spectrum of malignancies to construct a high-resolution pan-cancer bacteriome atlas. Initial profiling of α-diversity identified OV as harboring the highest bacterial complexity, followed sequentially by MB, LUAD, and BLCA (Figure 1D; Table S2). Across most cancer types, bacterial species richness demonstrated strong concordance with α-diversity indices, with the notable exception of MB (Figure S1D; Table S2). To probe the functional consequences of bacterial colonization within tumor tissues, we integrated bacterial diversity profiles with host transcriptomic landscapes. This analysis revealed significant associations between bacterial complexity and key oncogenic signaling programs. Specifically, tumors with high bacterial diversity exhibited marked enrichment in pathways driving cancer progression, such as apoptosis suppression and negative feedback regulation of the MAPK pathway^29^ (Figure 1E). Furthermore, these tumors displayed signatures associated with antibacterial responses (e.g., fatty acyl-CoA biosynthesis^29^) and mechanisms linked to bacterial internalization and tumor invasion, including NMDAR-mediated RAC1 activation^30–32^ (Figure 1E; Table S2). We next sought to determine whether these bacterial signatures were specific to the malignant niche. Cumulative genus distribution analysis identified BLCA and LUAD as harboring the most heterogeneous bacterial populations (Figure 1F; Table S2). Consistent with a potential role in tumorigenesis, bacterial diversity was significantly elevated in malignant tissues relative to matched normal-adjacent tissues (NATs, P < 0.01) (Figures 1G and 1H; Table S2). Finally, we utilized RNAscope-IF imaging to visualize the spatial distribution of these bacteria in situ. Our results confirmed the consistent presence of bacterial signals within both BLCA (Figure 1I) and LUAD (Figure 1J) specimens, with signals primarily localized within the tumor nest and exhibiting prominent co-localization with the cytoplasmic markers of tumor cells.

### Compositional divergence and tissue-specific signatures of the intratumoral bacteriome

To delineate the structural divergence of the intratumoral bacteriome, we performed comparative profiling across the pan-cancer cohort. Beta-diversity analysis using both Bray-Curtis and Jaccard distance metrics consistently demonstrated that gastrointestinal tumors harbored more similar bacterial communities within the same tumor type than across distinct tumor types (Figures 2A and S2A; Table S3), implicating shared anatomical or developmental origins in shaping their microbiome signatures. The distribution of family-level phylotypes revealed marked changes between the bacterial composition of the different tumor types (Figure 2B; Table S3). Consistent with prior studies, Pseudomonadota (Proteobacteria) emerged as the dominant phylum across multiple tumor types, including urothelial tract, liver, lung, ovarian, and prostate malignancies^33–36^ (Figure S2B; Table S3). Notably, Bacillota (Firmicutes) taxa, particularly the *Staphylococcaceae* family and *Staphylococcus* genus, were preferentially enriched in bladder tumors (Figure 2B and S2C; Table S3), while Actinomycetota represented a secondary but prevalent component in medulloblastoma and ovarian tumors (Figure S2B).

**Figure 2.**
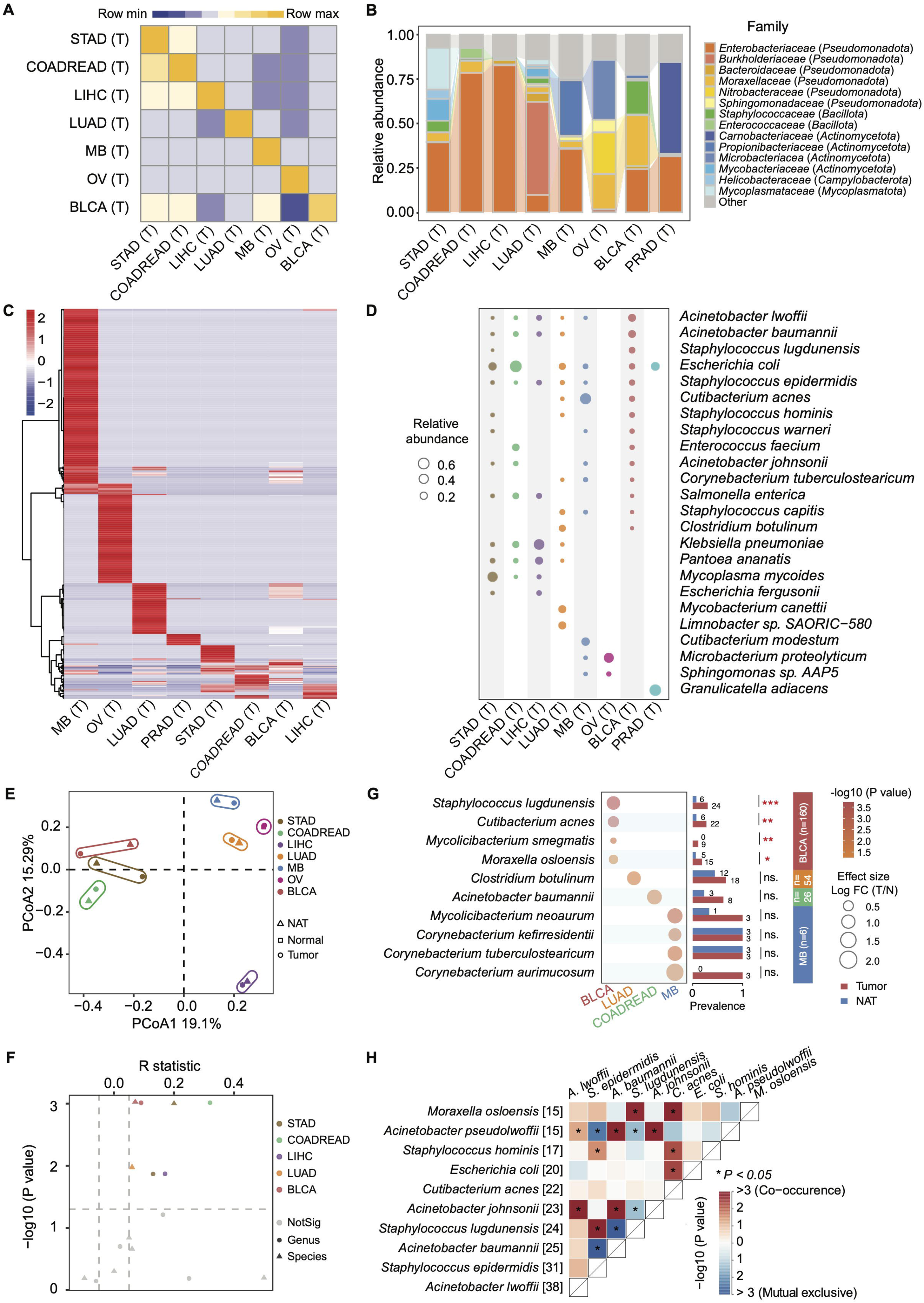
Diverse bacterial compositions and tumor associated signatures across cancer types. (A) Pairwise similarity heatmap illustrating the conservation of bacterial community structures across tumor types at the species level. (B) Relative abundance of dominant bacterial families within tumor tissues across the pan-cancer cohorts at the species level. (C) Clustered heatmap showing the relative abundance patterns of bacterial species (rows) across different cancer types (columns), revealing distinct niche specific bacterial signatures. (D) Bubble plot highlighting the relative abundance of selected bacterial species across diverse cancer types. The size of each dot represents the mean relative abundance within each cohort. (E) Principal Coordinates Analysis (PCoA) based on Bray Curtis dissimilarity at the species level, illustrating the distinct clustering of bacterial communities by cancer type. (F) Magnitude of bacterial community separation between tumor and NAT across the pan-cancer cohort. The scatter plot displays the ANOSIM R statistic (X axis) against the significance level (Y axis) at both the genus and species levels. The R statistic quantifies the degree of dissimilarity between bacterial communities in tumor versus NATs. (G) Comparative analysis of bacterial abundance and prevalence between tumor and matched normal adjacent tissue specimens at the species level. The bubble plot illustrates the natural log transformed fold change (Tumor versus NAT) and statistical significance of bacterial signatures as determined by MaAsLin2. Bar plots display the bacterial prevalence within each respective group. (H) Correlation matrix illustrating the co-occurrence (red) or mutual exclusivity (blue) between identified bacterial species (Fisher’s exact test). *, *P* < 0.05; **, *P* < 0.01; ***, *P* < 0.001. See also Figure S2, S3 and Table S3.

Species-level profiling unveiled distinct taxonomic partitioning across the cohort, with unsupervised clustering of the 415 most prevalent taxa identifying cancer-type-specific bacterial signatures (Figure 2C; Table S3). Notably, *S. lugdunensis* was selectively overrepresented in bladder tumors (Figure 2D; Table S3), a finding of interest as other members of the *Staphylococcus* genus have been previously implicated in breast cancer metastasis^37^. Other opportunistic pathogens also exhibited characteristic distributions, for instance, *Acinetobacter baumannii*, typically associated with nosocomial infections^38^, showed high abundance in bladder, lung, liver, and gastrointestinal malignancies (Figure 2D). Similarly, *Klebsiella pneumoniae*, a pathogen linked to pyogenic liver abscesses^39^, was enriched across lung, liver, and gastrointestinal tumors (Figure 2D). Furthermore, *Granulicatella adiacens*, a taxon previously associated with nasopharyngeal carcinoma risk^40^, was uniquely enriched in prostate adenocarcinoma (PRAD). Although unsupervised Principal Coordinate Analysis (PCoA) revealed that the global bacteriome across the pan-cancer cohort largely mirrored that of matched NATs (Figure 2E; Table S3), PERMANOVA identified significant compositional divergence β-diversity within bladder, gastrointestinal, liver, and lung malignancies (all *P* < 0.05) (Figures 2F, S2D, and S2E; Table S3). Taxonomic profiling at the species level corroborated these shifts, as average compositional distributions revealed pronounced alterations in bacterial proportions between tumor and NAT samples across diverse malignancies (Figure S3A; Table S3). Additionally, sample-specific taxonomic landscapes highlighted substantial inter-individual heterogeneity, reflecting the personalized microbial signatures inherent to individual tumor specimens (Figure S3B; Table S3). Consistent with these bacterial shifts, several taxa exhibited significant enrichment in tumor tissues relative to their matched NAT counterparts (Figure 2G; Table S3). Notably, *S. lugdunensis*, *Cutibacterium acnes*, and *Moraxella osloensis* were markedly overrepresented in BLCA, showing both increased prevalence and higher abundance compared with paired normal tissues (Figure 2G). Co-occurrence analysis further delineated the ecological organization of these enriched species, revealing a structured network of synergistic and antagonistic bacterial interactions (Figure 2H; Table S3). We observed a strong positive association between *S. lugdunensis* and *Staphylococcus epidermidis*, indicating their frequent co-existence within the malignant niche. Conversely, *S. lugdunensis* displayed a significant mutually exclusive relationship with *Acinetobacter baumannii*, suggesting competitive dynamics or distinct niche preferences among these tumor-associated bacteria (Figure 2H). Collectively, these findings establish a highly structured pan-cancer bacteriome landscape defined by profound compositional divergence and tissue-specific taxonomic signatures that differentiate malignant niches from their normal counterparts and potentially contribute to tumor progression.

### Machine learning identifies *S. lugdunensis* as a hallmark of bladder cancer

To interrogate the predictive potential of the intratumoral bacteriome, we developed gradient-boosting machine learning models leveraging normalized bacterial abundances (RPBM). These models exhibited robust discriminatory power across various taxonomic resolutions, achieving high classification performance for cancer types, stages, and tumor-versus-NAT differentiation (Figures 3A and S4; Table S4). Systematic evaluation of all prioritized features across these models identified *S. lugdunensis* exhibited pronounced tumor specificity in BLCA, demonstrating both significantly elevated abundance (FDR = 2.93 × 10^−2^) and 4-fold greater prevalence (FDR = 4.81 × 10^−3^) compared to NATs (Figures 3B and 3C; Table S4).

**Figure 3.**
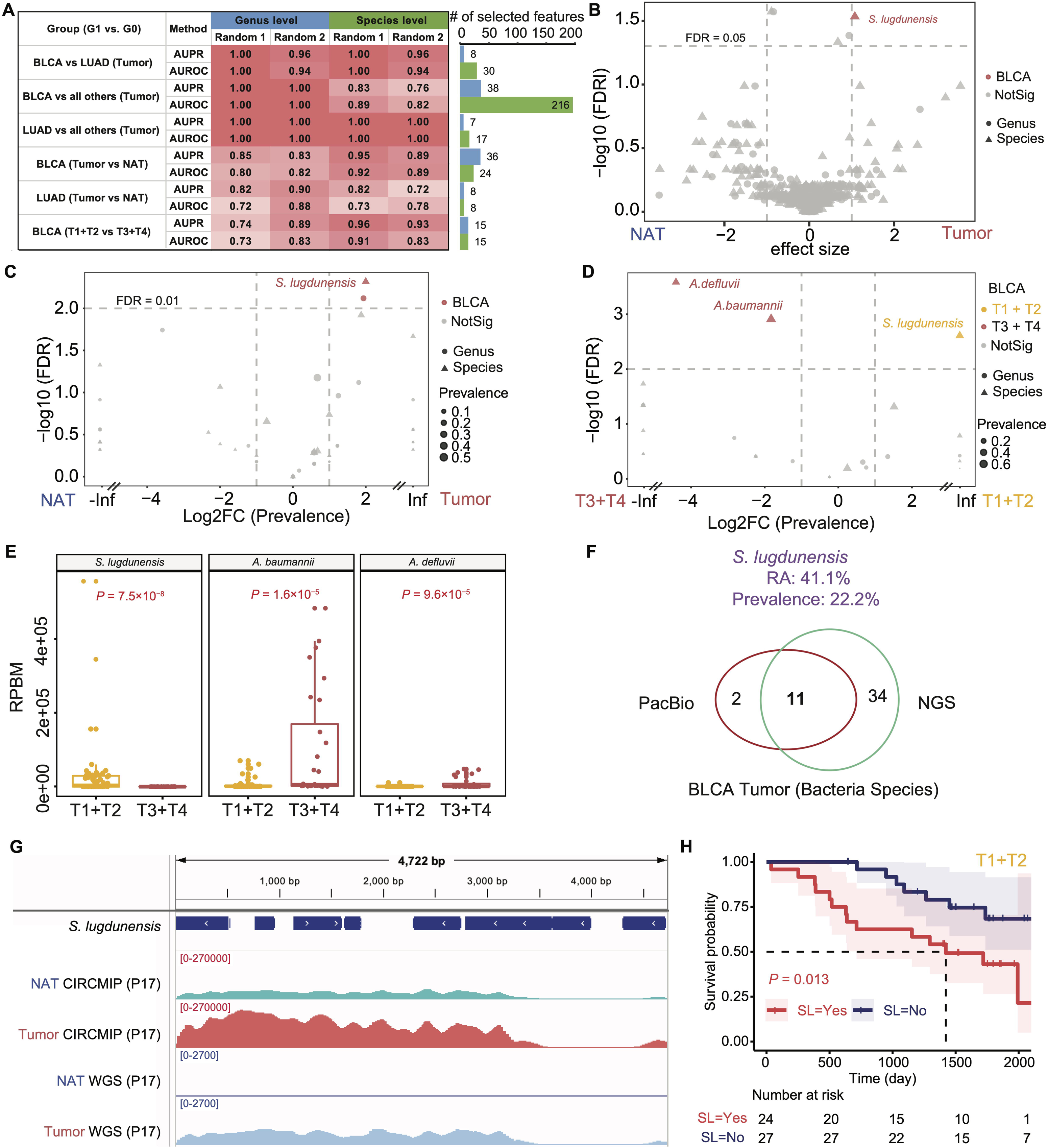
Performance of pan-cancer bacterial classification models and the clinical significance of *S. lugdunensis* in bladder cancer. (A) Performance evaluation of machine learning models for the classification of cancer types and tissue origins based on bacterial abundance. Heatmaps present the AUROC and AUPR scores at both genus and species levels, alongside the distribution of the number of selected bacterial features. (B) Volcano plot illustrating differentially abundant bacterial signatures between tumor and NATs across cancer types (MaAsLin2 algorithm using linear mixed-effects model adjusted for age and sex,). (C and D) Distribution of differences in bacterial prevalence between BLCA tumor and matched NAT (C) and between early (T1 plus T2) and late (T3 plus T4) clinical stages of BLCA (D). Statistical significance was assessed using the Mann Whitney U test. (E) Comparative analysis of bacterial abundance (RPBM) for *S. lugdunensis*, *Acinetobacter baumannii*, and *Acinetobacter defluvii* between early and late stage BLCA tumors. (F) Venn diagram illustrating the high consistency of bacterial species identification in BLCA tumor specimens between NGS and PacBio long read sequencing platforms. (G) Integrated Genomics Viewer (IGV) tracks comparing the genomic coverage of S. lugdunensis (NZ_CP084482.1) between CIRCMIP and standard WGS in a representative BLCA patient (P17). Note the significantly enhanced read depth provided by CIRCMIP in tumor (magenta) and NAT (cyan) tissues compared to conventional WGS (blue tracks). (H) Kaplan Meier survival analysis of early stage BLCA patients stratified by the presence (red) or absence (dark blue) of *S. lugdunensis* (SL) signatures within the tumor tissues (log rank test). See also Figure S4 and Table S4.

Notably, stage-specific interrogation revealed that the enrichment of *S. lugdunensis* was predominantly confined to early-stage bladder tumors (*P* = 7.5 × 10^−8^), whereas *Acinetobacter baumannii* (*P* = 1.6 × 10^−5^) and *Acinetobacter defluvii* (*P* = 9.6 × 10^−5^) predominated in late-stage malignancies (Figures 3D and 3E; Table S4). To substantiate these findings, we employed PacBio long-read sequencing as an orthogonal validation strategy, which confirmed the presence of *S. lugdunensis* in BLCA tumors (Figure 3F; Table S4). IGV visualization of *S. lugdunensis* genomic alignments corroborated the superior sensitivity of CIRCMIP over conventional WGS (Figure 1C). While WGS signals remained at baseline levels, CIRCMIP exhibited a ∼100-fold increase in dynamic range (scale 0-270,000 vs. 0-2,700), yielding robust and contiguous read coverage across the microbial locus in tumor tissues (Figure 3G). Notably, these signals were virtually undetectable in paired NATs, providing visual confirmation of CIRCMIP’s enhanced capacity to resolve low-abundance intra-tumoral bacterial signatures that are otherwise missed by standard metagenomic sequencing. Additionally, Kaplan-Meier survival analysis revealed its colonization was associated with markedly reduced overall survival in early-stage patients (*P* = 0.013) (Figure 3H; Table S4). Collectively, these results establish *S. lugdunensis* as a critical bacterial landmark in bladder cancer, whose presence not only distinguishes malignant status but also informs patient prognosis.

### *S. lugdunensis* drives bladder cancer progression via the LPC14:0-PPAR signaling axis

To examine the potential interplay between tumor-associated bacteria and the host immune microenvironment in bladder cancer (BLCA), we integrated the abundance profiles of prevalent bacterial species with immune cell infiltration estimated by CIBERSORT-based deconvolution of bulk transcriptomes. This analysis revealed that the abundance of *S. lugdunensis* was largely independent of inferred immune cell proportions (Figure 4A; Table S5), prompting us to investigate its direct effects on malignant cells. Co-culture assays demonstrated that *S. lugdunensis* significantly enhanced the proliferative capacity of T24 bladder cancer cells (Figure 4B; Table S5). Confocal imaging with HADA-labeled bacteria further visualized the intracellular localization of *S. lugdunensis* within tumor cells (Figure 4C), providing a spatial foundation for potential metabolic crosstalk between the bacteria and the malignant cell.

**Figure 4.**
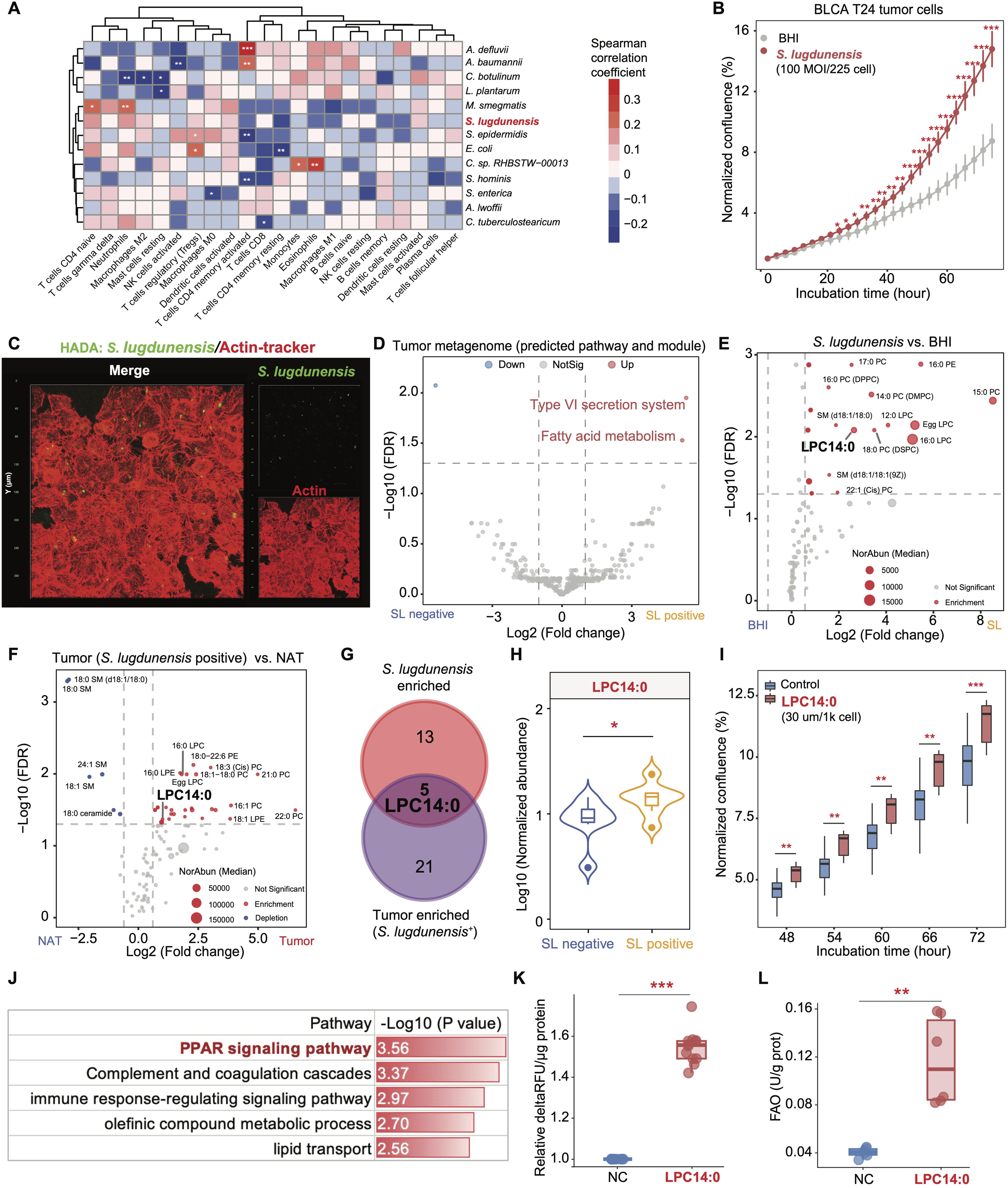
*S. lugdunensis* promotes bladder cancer progression via the LPC14:0-PPARδ axis. (A) Heatmap illustrating Spearman rank correlations between bacterial abundance and the infiltration proportions of diverse immune cell populations within the tumor microenvironment. (B) Real time proliferation curves of T24 BLCA cells following exposure to live *S. lugdunensis* at a multiplicity of infection of 100 to 1 compared to BHI medium control. Data represent the mean plus or minus standard error of the mean (SEM). (C) Representative confocal fluorescence imaging of T24 tumor cells following co culture with *S. lugdunensis*. Green fluorescent signals denote HADA labeled *S. lugdunensis*, which were observed localized within malignant cells visualized by red fluorescent actin tracker staining. (D) Volcano plot showing predicted bacterial metagenomic functional pathways and modules via PICRUSt2. (E and F) Volcano plots of differentially abundant lipid metabolites identified through targeted lipidomics, comparing *S. lugdunensis* lysates versus BHI medium (E) and *S. lugdunensis* positive tumors versus NATs (F). (G) Venn diagram depicting the intersection of enriched lipid species between bacterial lysates and tumor tissues, identifying LPC14:0 as a candidate effector molecule. (H) Normalized abundance of LPC14:0 in *S. lugdunensis* (SL) negative and positive tumor tissues. (I) Box plots showing the accelerated proliferation of T24 cells treated with exogenous LPC14:0 relative to vehicle control (six replicates/group). (J) Functional enrichment analysis of differentially expressed genes reveal that the PPAR signaling pathway was significantly enriched in the LPC14:0 treatment group. (K) Quantitative assessment of cellular fatty acid uptake in T24 cells following treatment with vehicle (NC) or 30 µM LPC14:0 for 18 hours. Data are expressed as relative background-subtracted fluorescence intensity (ΔRFU) normalized to total protein concentration (12 replicates/group). (L) Biochemical analysis of FAO activity in T24 cells under the same treatment conditions (six replicates/group). Two-tailed Student’s *t*-test: *, *P* < 0.05; **, *P* < 0.01; ***, *P* < 0.001; See also Figure S5 and Table S5.

Functional prediction via PICRUSt2 indicated an enrichment of fatty acid metabolism pathways in *S. lugdunensis* positive tumor tissues (Figure 4D; Table S5). To identify the molecular effectors of *S. lugdunensis* driving tumor progression, we conducted targeted lipidomic profiling across three comparative cohorts, including *S. lugdunensis* lysates versus BHI medium, colonization-positive tumor tissues versus matched NATs, and colonization-positive versus negative tumor tissues. Cross-referencing the differential signatures from the first two comparisons identified five metabolites that were consistently upregulated in both bacterial lysates and colonized tumor tissues relative to their respective controls (Figures 4E-G and S5; Table S5). Further evaluation of these five candidates within the clinical cohort of *S. lugdunensis* positive versus negative tumors confirmed the significant enrichment of LPC14:0 (Figure 4H), alongside LPC16:0 and Egg LPC (Figure S5B; Table S5), in colonized malignancies. Crucially, exogenous treatment with LPC14:0 was found to markedly enhance the proliferation of bladder tumor cells (Figures 4I, S5D, and S5F; Table S5).

Transcriptomic profiling of LPC14:0-treated cells revealed a robust activation of the PPAR signaling pathway (Figure 4J; Table S5). Consistent with these transcriptomic changes, functional assays demonstrated that LPC14:0 treatment significantly enhanced cellular fatty acid uptake (Figure 4K; Table S5). Furthermore, a marked increase in fatty acid beta oxidation (FAO) activity was observed in LPC14:0-treated cells (Figure 4L; Table S5). Collectively, these findings establish that *S. lugdunensis* facilitates bladder cancer progression by delivering LPC14:0, which in turn engages the host PPAR signaling axis to reprogram fatty acid metabolism and fuel malignant cell proliferation.

### The *S. lugdunensis*-LPC14:0 axis accelerates bladder cancer growth *in vivo*

To evaluate the pro-tumorigenic potential of *S. lugdunensis* and its identified metabolic effector LPC14:0 *in vivo*, we established a subcutaneous xenograft model using T24 bladder cancer cells in balb/c Null mice (Figure 5A). Following intragastric administration of *S. lugdunensis* and non-pathogenic *E. coli*, successful colonization of the bacteria within the tumor tissues was confirmed by re-isolation and plate culture from the excised specimens (Figure 5B; Table S6). Although both bacterial strains demonstrated the ability to persist in the tumor microenvironment, only *S. lugdunensis* significantly accelerated tumor expansion. Compared to the vehicle and *E. coli* controls, *S. lugdunensis* treatment led to markedly increased tumor volumes and terminal weights (Figures 5A-C; Table S6). Targeted lipidomic profiling validated the significant enrichment of LPC14:0 in *S. lugdunensis* relative to both the *E. coli* and vehicle controls (Figure 5D; Table S6). Consistent with the observed tumor expansion, immunohistochemical analysis of Ki67 expression further demonstrated that *S. lugdunensis* colonization substantially augmented the proliferative capacity of malignant cells (Figures 5E and 5F; Table S6). Furthermore, *S. lugdunensis* treatment was associated with a dramatic elevation in FAO activity within the tumor mass, establishing a link between bacterial colonization and enhanced lipid catabolism (Figure 5G; Table S6).

**Figure 5.**
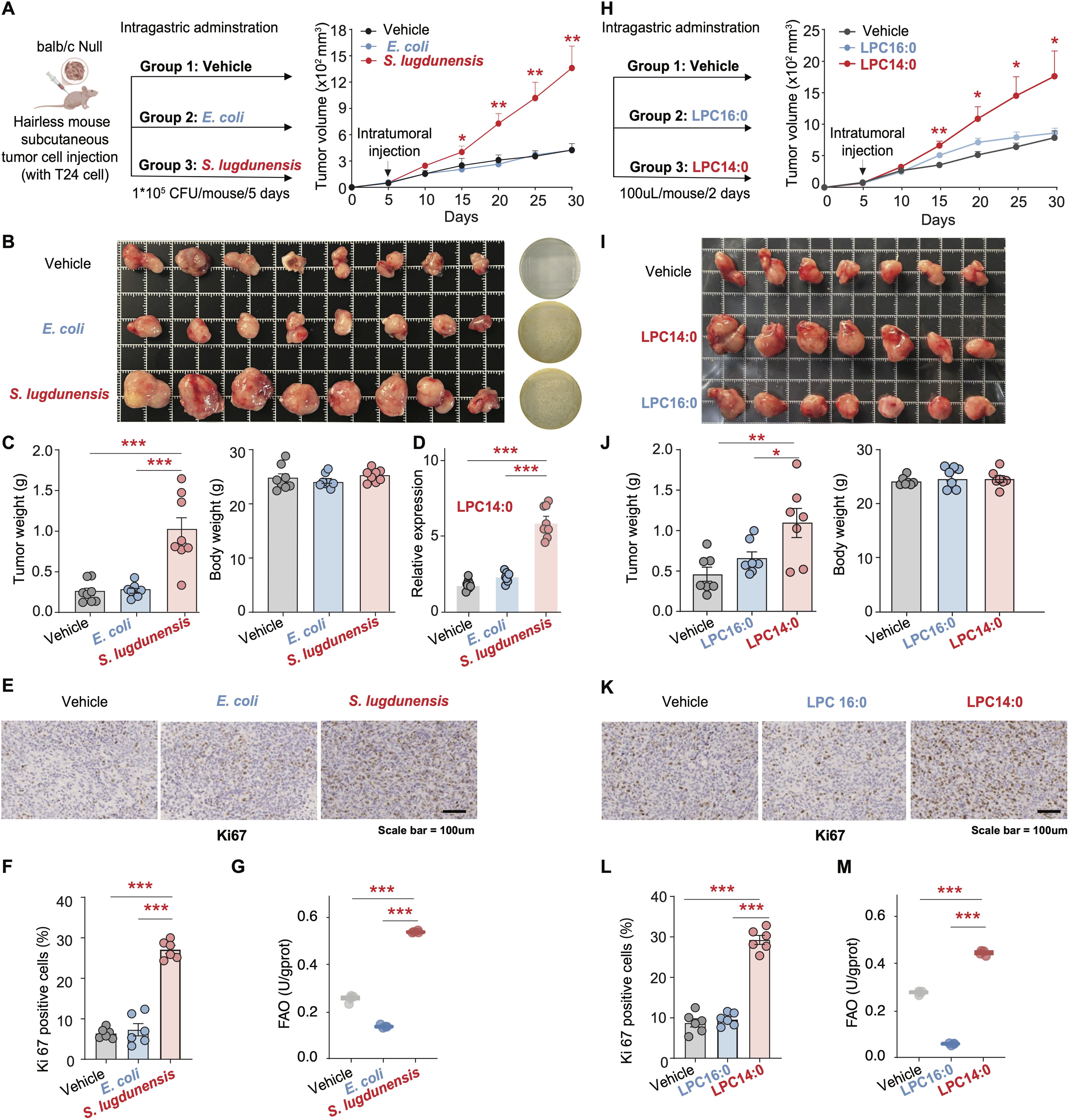
*S. lugdunensis* and its metabolite LPC14:0 promote bladder cancer progression in vivo. (A) Schematic representation of the experimental workflow in balb/c Null hairless mice. T24 bladder cancer cells were subcutaneously injected, and mice were subsequently treated with vehicle, *Escherichia coli*, or *S. lugdunensis* at the fifth days. Tumor volume was monitored over 30 days, showing that *S. lugdunensis* significantly accelerated tumor growth. (B) Representative tumor images of resected tumors at day 30 and corresponding bacterial colony forming units recovered from the tumor tissues, confirming successful bacterial colonization. (C) Quantitative analysis of terminal tumor weight and longitudinal mouse body weight across the three experimental groups (N=8/group). (D) Difference of relative expression of LPC14:0 across the three experimental groups. (E) Representative immunohistochemistry (IHC) images of Ki67 staining in tumor sections. Scale bars represent 100 microns. (F) The bar graph shows the quantification of Ki67 positive cells, indicating enhanced cellular proliferation in the *S. lugdunensis* group. (G) Biochemical analysis of FAO activity in mice tumor under the *S. lugdunensis* conditions (N=4/group). (H) Experimental design for the exogenous administration of the LPC14:0. Mice harboring T24 tumors received intratumoral injections of vehicle or LPC14:0. Growth curves demonstrated that LPC14:0 significantly enhanced tumor expansion. (I) Gross morphology of tumors harvested from the vehicle and LPC14:0 treated groups at the study endpoint. (J) Statistical comparison of final tumor weights and mouse body weights between the vehicle and LPC14:0 groups (N=7/group). (K-L) IHC analysis of Ki67 expression in tumor tissues from mice treated with LPC14:0. Scale bars represent 100 microns. The percentage of Ki67 positive cells was significantly higher in the LPC14:0 group, confirming the pro proliferative effect of the metabolite. (M) Biochemical analysis of FAO activity in mice tumor under the LPC14:0 treatment conditions (N=4/group). Two-tailed Student’s *t*-test: *, *P* < 0.05; **, *P* < 0.01; ***, *P* < 0.001; See also Figure S5 and Table S6.

We further interrogated whether the downstream metabolite LPC14:0 could recapitulate the tumorigenic phenotype observed with bacterial colonization, using LPC16:0 as a structural analog control to assess specificity. Consistent with the effects of *S. lugdunensis*, administration of LPC14:0 resulted in significantly accelerated tumor growth and a higher final tumor burden (Figures 5H-J; Table S6). In contrast, treatment with LPC16:0 had a negligible impact on tumor progression, with the overall tumor burden remaining comparable to that of the vehicle control (Figures 5H-J). This induction by LPC14:0 was accompanied by a robust increase in the proportion of Ki67-positive cells (Figures 5K-L; Table S6). Notably, functional assays confirmed that LPC14:0, but not its analog LPC16:0, specifically stimulated FAO activity in the tumor tissues (Figure 5M; Table S6). Collectively, these *in vivo* findings provide definitive evidence that the *S. lugdunensis*-LPC14:0 axis functions as a potent driver of bladder cancer progression.

### LPC14:0–PPAR**δ** direct activation orchestrates a pro-tumorigenic fatty acid metabolism

To elucidate the molecular mechanism underlying the pro-tumorigenic effects of LPC14:0, we first employed a chemical proteomics approach using a bifunctional photo-crosslinking probe (*LPC14:0; Figure 6A). In situ fluorescence imaging in T24 cells revealed robust labeling by the LPC14:0 probe, which was specifically outcompeted by excess native LPC14:0 (100 μM) (Figure 6B). This competitive binding demonstrates that the bifunctional probe recapitulates the binding behavior of its natural counterpart, providing a validated surrogate for the subsequent proteomic identification of endogenous targets. Affinity purification followed by LC-MS/MS analysis identified 31 potential protein targets of LPC14:0 in T24 cells. Given the activation of the PPAR signaling pathway observed in our previous transcriptomic analysis (Figure 4J), we cross-referenced these MS-identified candidates with components of this pathway. This integrated analysis revealed PPARδ as a prominent binding target of LPC14:0 (Figure 6C; Table S7), supporting the notion that LPC14:0 acts as a direct ligand for this nuclear receptor. To validate the functional impact of the *S. lugdunensis*–LPC14:0 axis *in vivo*, we performed transcriptomic profiling on xenograft tumor tissues. Gene Set Enrichment Analysis (GSEA) revealed that both *S. lugdunensis* colonization and LPC14:0 treatment significantly enriched pathways associated with systemic lipid homeostasis and catabolism, including “Fatty acids”, “Free fatty acid receptors”, and “Ldl clearance” (Figures 6D-E; Table S7). Specifically, volcano plot analysis showed consistent upregulation of key fatty acid metabolism genes (*CD36*, *FABP4*, *ACOX2*) in response to *S. lugdunensis* and LPC14:0, in contrast to the lack of significant induction by LPC16:0 treatment (Figures 6F-G; Table S7). These results strongly suggest that the *S. lugdunensis*-derived LPC14:0 triggers a concerted metabolic reprogramming within the tumor.

**Figure 6.**
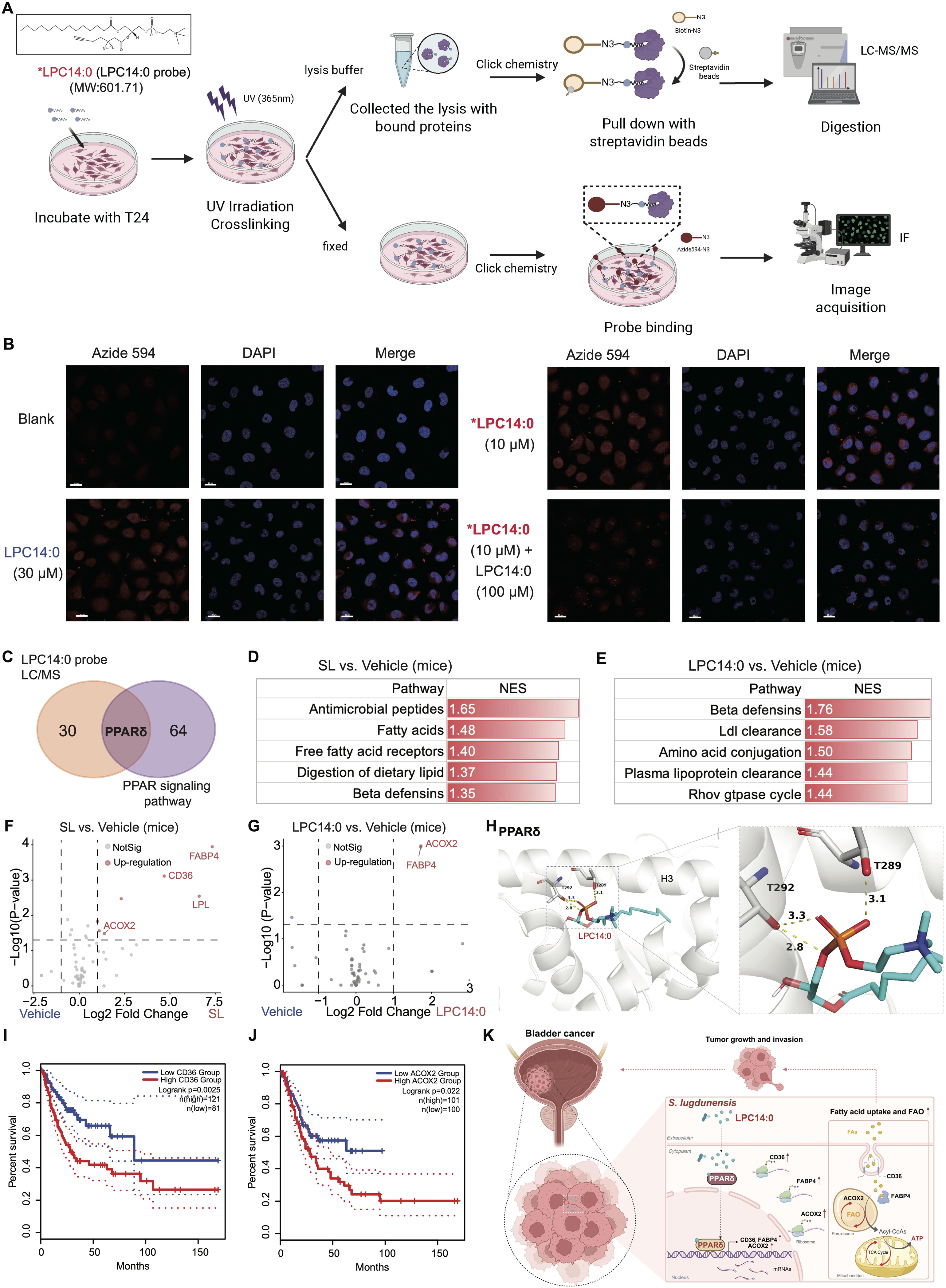
LPC14:0 directly activates host PPARδ to reprogram fatty acid metabolism and drive bladder cancer progression. (A) Schematic representation of the chemical proteomics workflow utilizing a bifunctional photo-crosslinking probe (*LPC14:0) for target protein capture, visualization, and identification in T24 cells. (B) *In situ* fluorescence imaging of T24 cells following incubation with *LPC14:0 and bioorthogonal conjugation with Azide 594 (red). Specific binding is demonstrated by competitive binding of the probe signal with 10-fold excess native LPC14:0 (100 µM). Nuclei were counterstained with DAPI (blue). Scale bars, 20 µm. (C) Venn diagram showing the intersection of MS-identified LPC14:0-interacting proteins with the PPAR signaling pathway, pinpointing PPARδ as a primary candidate. (D, E) GSEA pathway enrichment analysis in tumor tissues from mice treated with *S. lugdunensis* (D) or LPC14:0 (E). Significant enrichment of pathways is ranked by Normalized Enrichment Scores (NES). (F, G) Volcano plots representing differentially expressed genes in *S. lugdunensis*-treated (F) and LPC14:0-treated (G) tumors. (H) Molecular docking model illustrating the binding configuration of LPC14:0 (green sticks) within the active pocket of PPARδ (gray ribbons) (Vina score=-6.5; Cavity volume= 408 Å). The inset provides a detailed view of stable hydrogen bonds formed with amino acid residues Thr292 and Thr289 (yellow dashed lines; distances <3.5 Å). Atoms are color coded as follows: red, oxygen; orange, phosphorus; blue, nitrogen. (I, J) Kaplan-Meier curves of overall survival in bladder cancer patients (TCGA cohort, via GEPIA2) stratified by *CD36* (I) or *ACOX2* (J) expression levels. Statistical significance was determined by the log-rank test. (K) A proposed working model: *S. lugdunensis* secretes LPC14:0 into the tumor tissue, where it directly engages and activates host PPARδ. This activation triggers the transcriptional upregulation of fatty acid transporters and oxidases, fueling malignant progression through enhanced lipid uptake and bioenergetic demand. *, *P* < 0.05; **, *P* < 0.01; ***, *P* < 0.001; See also Figure S5 and Table S7.

To gain structural insights into the interaction between LPC14:0 and its putative target, molecular docking was performed. The results demonstrated that LPC14:0 fits precisely into the ligand-binding domain of PPARδ, forming stable hydrogen bonds with critical residues, including Thr292 and Thr289 (Figure 6H). This high-affinity binding provides a structural basis for the activation of PPARδ and the subsequent transcriptional induction of its downstream metabolic targets. Consistent with their pro-tumorigenic roles, increased expression of the PPARδ target genes *CD36* and *ACOX2* predicted significantly worse survival in bladder cancer patients, confirming the prognostic impact of the *S. lugdunensis*–LPC14:0–PPARδ axis (Figures 6I-J; Table S7). Collectively, these findings delineate a comprehensive mechanistic model (Figure 6K): *S. lugdunensis* infiltrates the bladder tumor tissues and secretes LPC14:0, which directly binds to and activates the host nuclear receptor PPARδ. This activation drives the transcriptional up-regulation of *CD36*, *FABP4*, and *ACOX2*, thereby enhancing cellular fatty acid uptake, intracellular transport, and peroxisomal β-oxidation. This metabolic shift not only provides essential energy (ATP) through FAO but also yields structural precursors for membrane biogenesis, ultimately fueling the rapid proliferation and progression of bladder cancer.

## DISCUSSION

The tumor-resident microbiome represents an emerging hallmark of cancer, yet progress in this field has been constrained by formidable technical obstacles, including extremely low bacterial biomass, pervasive contamination risks, and the overwhelming abundance of host DNA. In this study, we developed CIRCMIP, a circular DNA-based enrichment and analytical framework that overcomes these limitations by exploiting the fundamental topological distinction between bacterial and eukaryotic genomes. By selectively capturing circular bacterial DNA, CIRCMIP achieved a more than 100-fold increase in bacterial read detection compared to conventional whole-genome shotgun metagenomics, enabling high-confidence profiling of the intratumoral bacteriome across a pan-cancer cohort of eight malignancies.

Application of CIRCMIP revealed a highly structured landscape of tumor-associated bacteria characterized by cancer type-specific taxonomic signatures. Consistent with previous reports^33–36^, Pseudomonadota constituted the dominant phylum across most malignancies. Notably, we identified marked enrichment of Bacillota, particularly the Staphylococcaceae family and *Staphylococcus* genus, in bladder tumors. Crucially, this observation attained statistical significance solely through the enhanced sensitivity of our circular DNA-based approach. Species-level resolution further pinpointed *S. lugdunensis* as selectively overrepresented in bladder cancer, exhibiting both significantly elevated abundance and fourfold greater prevalence in malignant tissues compared to matched normal-adjacent specimens. These observations align with recent evidence implicating other Staphylococcus species in breast cancer metastasis^37^ and suggest that staphylococcal taxa may harbor broader oncogenic potential across epithelial malignancies than previously appreciated.

Functionally, our findings establish *S. lugdunensis* and its primary metabolic effector, LPC14:0, as potent drivers of bladder cancer progression. Both *in vitro* co-culture assays and *in vivo* xenograft models demonstrated that *S. lugdunensis* colonization significantly accelerates malignant cell proliferation and tumor expansion. Notably, this oncogenic effect is highly specific to the 14:0 acyl chain, as the structural analog LPC16:0 failed to elicit comparable tumor growth, highlighting the unique bioactivity of LPC14:0. This pro-tumorigenic phenotype is underpinned by a profound metabolic reprogramming. Functional assays demonstrated that LPC14:0 treatment facilitates fatty acid uptake and β-oxidation, thereby fulfilling the massive bioenergetic and biosynthetic requirements of rapidly proliferating tumor cells.

Mechanistically, *S. lugdunensis*-derived LPC14:0 directly activates the host PPARδ axis to fuel tumor metabolic reprogramming. The direct binding of LPC14:0 to PPARδ was confirmed through chemical proteomic profiling with a photo-crosslinking probe, while molecular docking elucidated the structural basis of this interaction, highlighting critical hydrogen bonds with Thr292 and Thr289. PPARδ activation subsequently upregulates downstream targets including *CD36*, *FABP4*, and *ACOX2*, driving a metabolic reprogramming program that sustains malignant cell expansion. The convergence of evidence across orthogonal experimental systems, spanning bacterial-cancer cell co-culture, targeted lipidomic profiling, transcriptomic analysis, establishes the LPC14:0–PPARδ axis as a pivotal effector mechanism, further supported by *in vivo* xenograft modeling. These findings extend the emerging paradigm of microbial metabolite-driven oncogenesis^15,24^ and establish lysophospholipids as a class of bacterially derived signaling molecules with tumor-promoting capacity.

The clinical implications of our findings are multifaceted. First, the robust diagnostic performance of CIRCMIP-derived bacterial biomarkers (AUC = 0.92) suggests that intratumoral bacterial signatures may serve as pivotal adjuncts to conventional tissue-based diagnostics for refining patient stratification and clinical decision making in bladder cancer. Second, the prognostic association between *S. lugdunensis* colonization and reduced overall survival in early-stage patients identifies this bacterium as a potential stratification biomarker for identifying high-risk individuals who may benefit from more intensive surveillance or adjunctive therapies. Third, the delineation of the LPC14:0–PPARδ axis reveals actionable therapeutic targets. Pharmacological PPARδ antagonists, already developed for metabolic disorders, could potentiallybe repurposed to disrupt this tumor-promoting pathway, while antibiotics targeting *S. lugdunensis* might offer a strategy for early intervention. However, the latter approach must be approached cautiously given the risk of disrupting protective commensal communities and the potential for off-target effects on the gut microbiome^4–6^.

Looking forward, the CIRCMIP framework establishes a new paradigm for intratumoral bacteriome research that can be extended in several directions. Integration with single-cell RNA sequencing technologies could resolve bacterial heterogeneity at the cellular level and identify host cell subpopulations preferentially colonized by specific taxa^41^. Spatial transcriptomics coupled with bacterial probe-based detection would enable mapping of microbe–host interaction niches within the tumor architecture. Functional metagenomic profiling of bacterial isolates from patient tumors could uncover additional metabolite-mediated mechanisms of host modulation. Moreover, application of CIRCMIP to longitudinal cohorts with detailed treatment histories may reveal whether intratumoral bacterial composition influences therapeutic responses to immunotherapy, chemotherapy, or radiation—analogous to the established role of the gut microbiome in modulating anti-PD-1 efficacy^4–6^.

### Limitations of the study

Some limitations of this study should be acknowledged. First, while our subcutaneous xenograft models demonstrated the pro-tumorigenic effects of *S. lugdunensis* and LPC14:0, orthotopic bladder cancer models incorporating a functional immune system would better recapitulate the native tumor microenvironment and enable assessment of immune-modulatory effects. Second, our cohort, while comprehensive across eight cancer types, included relatively limited numbers of certain malignancies (e.g., medulloblastoma, prostate adenocarcinoma), and validation in larger, independent cohorts will be essential to confirm the generalizability of our findings.

## RESOURCE AVAILABILITY

### Lead contact

Further information and requests for resources and reagents should be directed to and will be fulfilled by the lead contact, Fengbiao Mao.

### Materials availability

This study did not generate new unique reagents.

### Data and code availability

A detailed overview of data accessibility, including primary data sources and specific accession identifiers for all high throughput sequencing datasets, was provided in Table S1. The raw circular DNA enrichment data, whole genome sequencing (WGS) profiles, and RNA sequencing (RNA seq) datasets, along with associated and clinical metadata, have been submitted to the Genome Sequence Archive (GSA) with ID HRA017734, HRA017801, CRA041086 and CRA041087. All other data supporting the findings of this study are available within Supplementary Tables S2 to S7. Any additional information required to reanalyze the data reported in this paper is available from the lead contact upon request.

## ACKNOWLEDGMENTS

This work was supported by the National Natural Science Foundation of China [Grant No. 32470835, No. 32170493], the Beijing Natural Science Foundation (Grant No. 7242169, No. L248056), the Clinical Medicine Plus X - Young Scholars Project, Peking University, the Fundamental Research Funds for the Central Universities (Grant No. PKU2024LCXQ044).

## AUTHOR CONTRIBUTIONS

FB Mao, PC Wang and T Wang conceived and designed the study. FB Mao, PC Wang, T Wang, XL Zhao and M Zhao designed the experiments. GR Xiao and DY Kong designed and synthesized the LPC14:0 probe. PC Wang, M Zhao and HY Zhao performed the experiments. W Lv, F Zhang, CH Li, and CH Lin collected the clinical samples. T Wang performed the analysis and wrote the original draft of the manuscript. FB Mao, PC Wang and T Wang revised and edited the paper. All authors improved the manuscript and approved the submission.

## SUPPLEMENTAL INFORMATION

**Document S1. Figures S1-S5 and Table S1 - S7**

**Table S1. Metadata for donors, and samples included in this study, related to Figure 1**.

**Table S2. Taxonomic abundance, diversity metrics, and pan-cancer landscape data of the intratumoral bacteriome, related to Figures 1, S1, and S2.**

**Table S3. Differentially enriched bacterial signatures and species-level taxonomic composition across diverse cancer types, related to Figures 2 and S3.**

**Table S4. Machine learning model performance and clinical association analysis of *S. lugdunensis* in bladder cancer, related to Figure 3 and S4.**

**Table S5. Transcriptomic GSEA results and metabolic characterization of *S. lugdunensis* and its lipid derivatives, related to Figure 4 and S5.**

**Table S6. *In vivo* xenograft tumor growth parameters, Ki67 quantification, and metabolic phenotyping data, related to Figure 5**.

**Table S7. Targeted proteomics and clinical survival analysis of the LPC14:0-PPAR**δ **axis, related to Figure 6**.

**Figure S1. High resolution profiling and taxonomic refinement of the pan-cancer intratumoral bacteriome.** (A and B) Relative abundance of diverse genomic components, including fungi, archaea, viruses, bacteria, plasmids, and mitochondria. Data are presented for tumor specimens (A) across eight cancer types and pan-cancer NATs (B). (C) The five-stage taxonomic filtration module within the CIRCMIP framework for high confidence bacterial species identification. This hierarchical quality control process was implemented within the CIRCMIP computational framework to progressively refine bacterial signatures across 312 samples from 8 cancer types. From an initial pool of 7,283 bacterial species, stringency was achieved through five integrated filters. Filter 1 excluded laboratory and library contaminants. Filter 2 performed sample level signal denoising. Filter 3 utilized negative control cell line data to establish background thresholds. Filter 4 implemented healthy tissue reference-based filtering to prioritize tumor enriched signals. Finally, Filter 5 removed low abundance taxa to minimize stochastic noise. This integrated pipeline enabled the robust identification of 732 high confidence bacterial species within the tumor tissues. (D) Quantitative distribution of the number of identified bacterial species per sample across the eight solid tumor types. Each dot represents an individual specimen, illustrating the variations in bacterial richness across different clinical cohorts.

**Figure S2. Bacterial community structure and beta diversity across diverse solid tumor types.** (A) Jaccard similarity heatmap illustrating the degree of overlap in bacterial species composition across solid tumor types at the species level. The color gradient represents row normalized similarity scores, reflecting the conservation of bacterial presence between different malignancies. (B and C) Distribution of the relative abundance of bacterial phyla (B) and genera (C) across the cancer cohorts. (D and E) Principal Coordinates Analysis (PCoA) characterizing bacterial beta diversity using quantitative Bray Curtis dissimilarity (D) and qualitative Jaccard similarity (E) distance metrics. Individual samples and corresponding ellipses are colored by tissue origin, including tumor specimens (red) and NATs (blue). Statistical significance and the effect size of the compositional differences were determined via permutational multivariate analysis of variance (PERMANOVA), with the resulting R values and P values annotated for each cancer type.

**Figure S3. Taxonomic composition and inter individual heterogeneity of the intratumoral bacteriome at the species level.** (A) Stacked bar plots illustrating the mean relative abundance of the top bacterial species across solid tumor types, comparing tumor specimens and NATs. (B) Detailed species level abundance profiles for individual patient samples within each cancer cohort. Each vertical bar corresponds to a single specimen, revealing the significant inter individual heterogeneity of the bacterial community structure within the tumor tissues.

**Figure S4. Performance of bacterial abundance-based machine learning models for BLCA classification.** Precision Recall (PR) and Receiver Operating Characteristic (ROC) curves evaluating the performance of classifiers in distinguishing BLCA tumor specimens from matched NATs. The models were trained and validated at both the genus level (left columns) and species level (right columns). (B) Diagnostic performance of bacterial signatures in discriminating early stage (T1 plus T2) from late stage (T3 plus T4) BLCA tumors. The color gradient on the curves represents the variation in prediction probability thresholds.

**Figure S5. Metabolic characterization of S. lugdunensis and the impact of its lipid derivatives on bladder cancer cell proliferation.** (A) Partial least squares discriminant analysis (PLS DA) illustrating the divergent lipidomic profiles between *S. lugdunensis* (SL) lysates (red circles) and BHI medium controls (green circles). The axes represent the percentage of total variance explained by the first two components. (B) Log 10 normalized abundance of candidate lipid species, including LPC16:0, Egg LPC, PC14:0, and PC18:0, in clinical bladder cancer specimens stratified by the presence (SL positive) or absence (SL negative) of *S. lugdunensis* signatures. (C) Quantitative assessment of specific lipid metabolites in *S. lugdunensis* lysates compared to BHI medium. Bar graphs represent the normalized abundance of LPC16:0, Egg LPC, LPC14:0, PC14:0, and PC18:0, showing significant enrichment within the bacterial fraction. (D to F) Real time monitoring of T24 BLCA cell proliferation following treatment with exogenous lipids: LPC16:0 (D), Egg LPC (E), and PC14:0 (F). Confluence percentages were recorded over a period of 72 hours and normalized to initial measurements at 0 hours. Data are presented as the mean plus or minus standard error of the mean (SEM). Statistical significance was determined using the Student’s t test, where asterisks indicate *P* values below 0.05.

## STAR⍰METHODS

### KEY RESOURCES TABLE

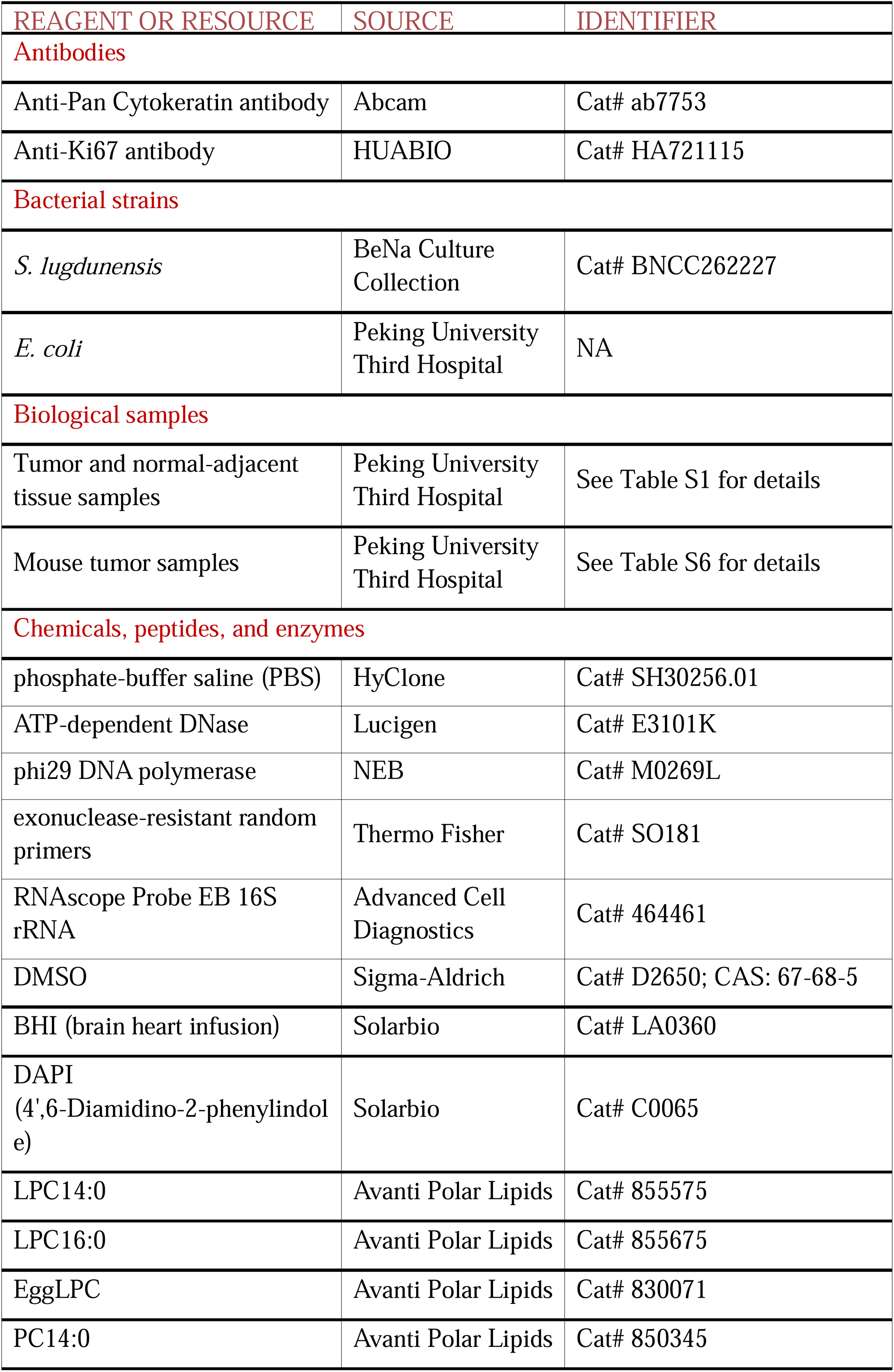
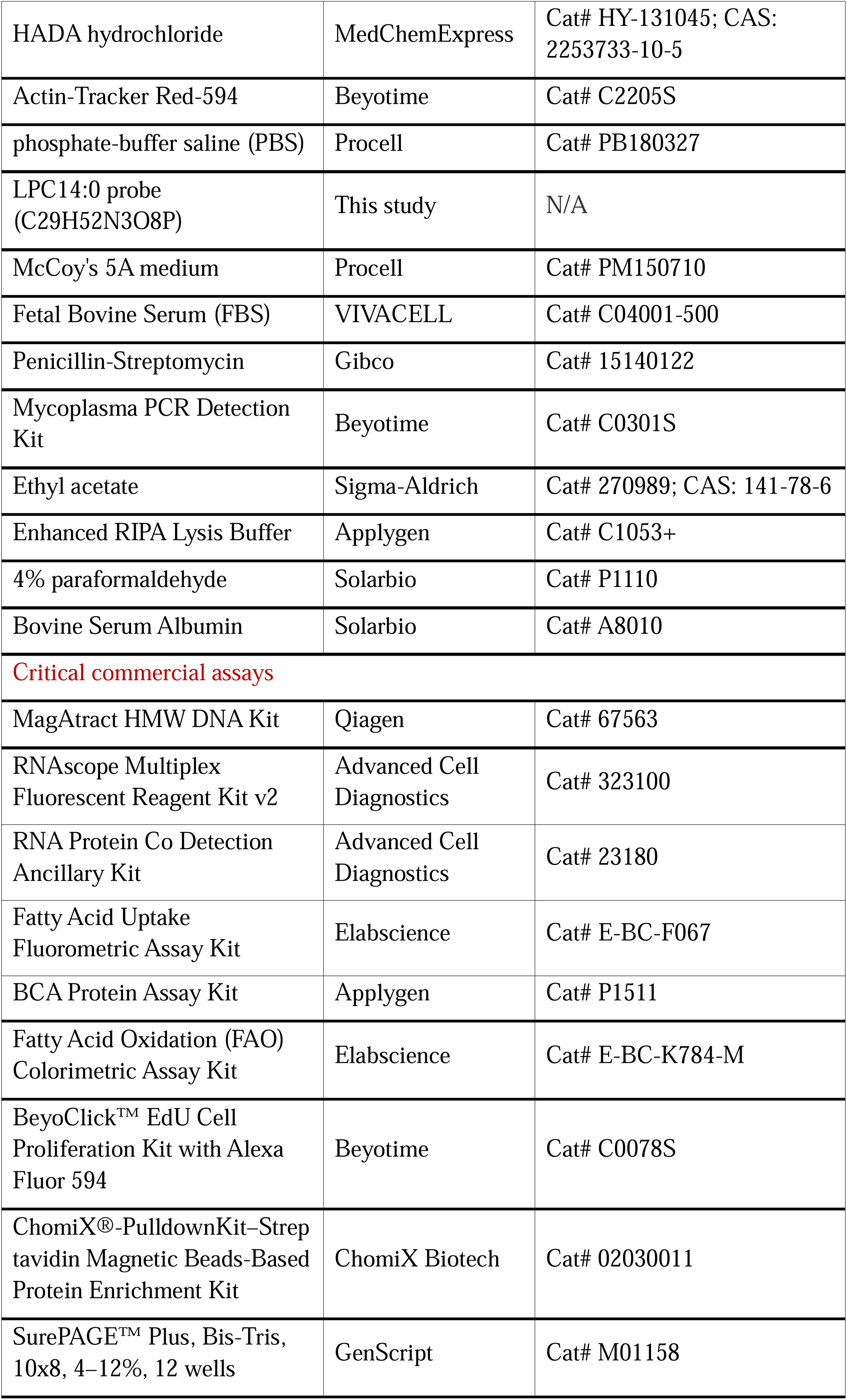
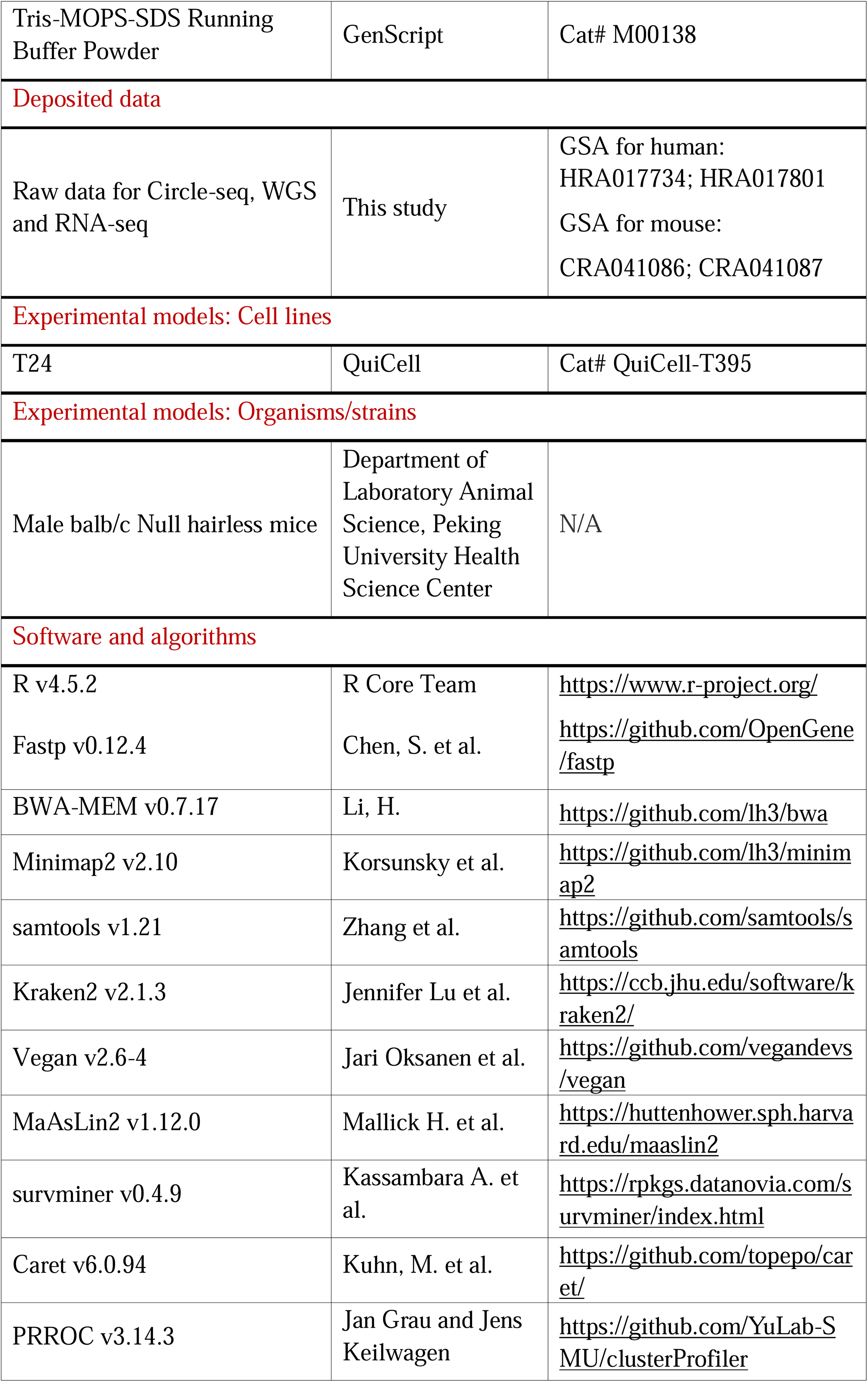
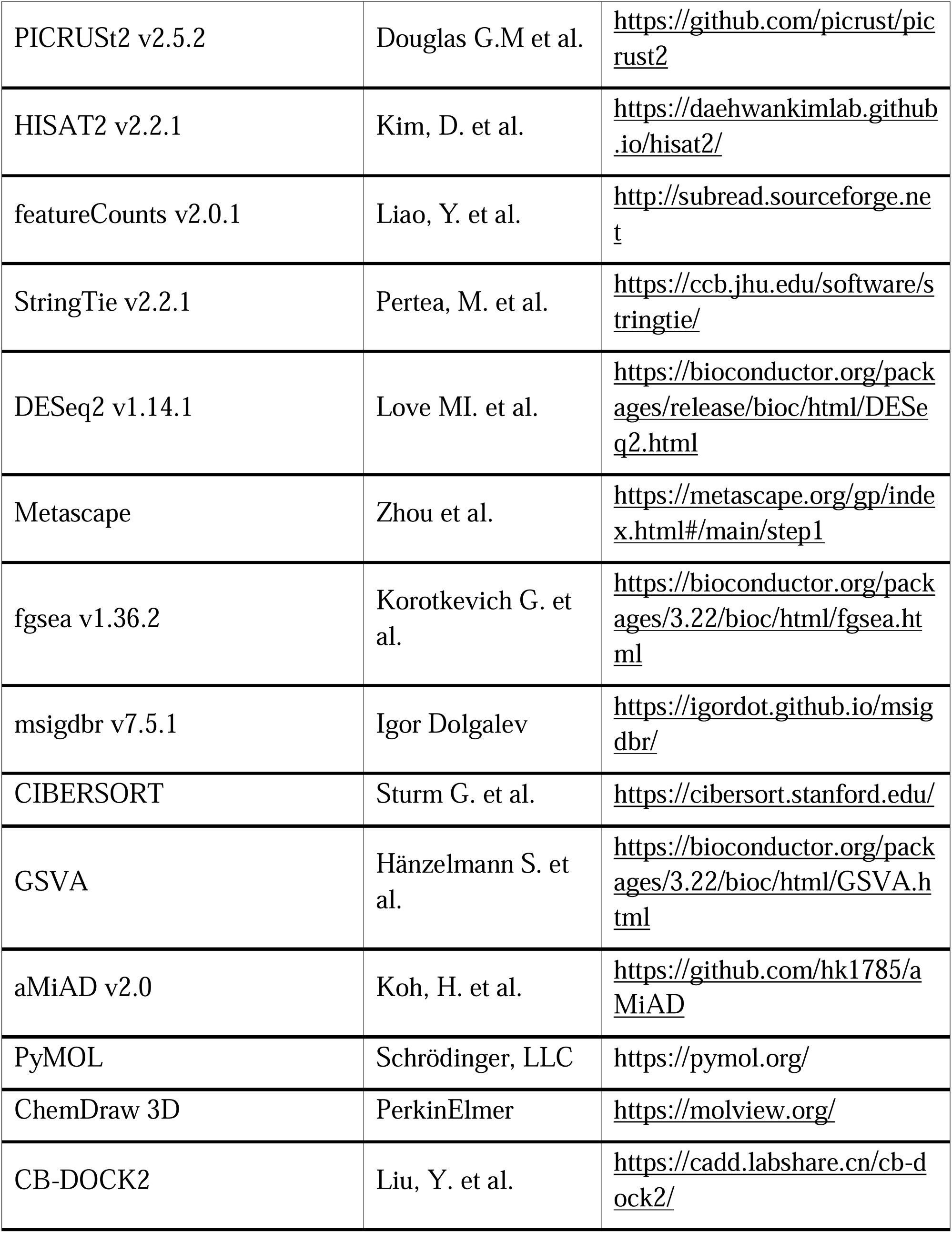

### EXPERIMENTAL MODEL AND STUDY PARTICIPANT DETAILS

#### Clinical sample

This comprehensive study analyzed 312 biospecimens from 156 cancer patients and 5 healthy volunteers across eight cancer types, including 148 matched tumor-normal adjacent tissue (NAT) pairs (80 BLCA, 13 COADREAD, 10 LIHC, 27 LUAD, 13 STAD, 3 MB, 2 PRAD) along with 11 unmatched OV and 5 normal ovary controls (complete sample characteristics in Table S1)^42–46^. For comparative analysis, we performed: (i) whole-genome sequencing (WGS), and (ii) Circle-seq in an independent BLCA cohort (Table S1).

#### Cell line

The human bladder cancer cell line T24 was obtained from QuiCell (Cat# QuiCell-T395). T24 cells were cultured in McCoy’s 5A medium (Procell, Cat# PM150710) supplemented with 10% (v/v) fetal bovine serum (FBS; VIVACELL, Cat# C04001-500) and 1% (v/v) penicillin-streptomycin (Gibco, Cat# 15140122). All cells were maintained in a humidified incubator at 37°C with a 5% CO_2_ atmosphere. The cell line was routinely tested and confirmed to be negative for mycoplasma contamination (Beyotime, Cat# C0301S). Cell line identity was authenticated by Short Tandem Repeat (STR) profiling.

#### Animal models

Male balb/c Null hairless mice (6 to 8 weeks old) were utilized for all animal experiments. Mice were given a normal chow diet that meets national standards GB14924.3-2010 in all animal experiments. Mice had free access to water and food and were housed in a 12□h light/12□h dark cycle, with the temperature kept at 21-24□°C and humidity at 40-70%. All animal experimental were approved by the Animal Care and Use Committee of Peking University Health Science Center according to the national legislation for animal care (Animal Protocol Approval No.: DLASBE0547). To establish the bladder cancer xenograft model, one million T24 cells (Cat# QuiCell-T395) were subcutaneously injected into the right flank of each mouse. Mice were maintained under specific pathogen free conditions, and all procedures were performed in accordance with institutional ethical guidelines for animal care.

### METHOD DETAILS

#### Library preparation and circular DNA sequencing

Samples were washed with phosphate-buffer saline (PBS, HyClone, Cat# SH30256.01) immediately following surgical resection to remove blood. Samples (∼30 mg) were cut into small pieces, and total genomic DNA was extracted using the MagAtract HMW DNA Kit (Qiagen, Cat# 67563).

To isolate circular DNA, linear DNA was digested using plasmid safe ATP-dependent DNase (Lucigen, Cat# E3101K) in a 50 μL reaction at 37°C for 48 hours. To ensure exhaustive digestion, the reaction was replenished every 24 hours with 1 µL of ATP, and 1 µL of plasmid safe ATP-dependent DNase, according to the manufacturer’s agreement. The complete removal of linear DNA was validated by PCR amplification of the internal control gene *Cox5b* (F: 5’-GGGCACCATTTTCCTTGATCAT-3’; R: 5’-AGTCGCCTGCTCTTCATCAG-3’). Samples verified as *Cox5b*-negative were subjected to Rolling Circle Amplification (RCA) using phi29 polymerase (NEB, Cat# M0269L) and exonuclease-resistant random primers (Thermo Fisher, SO181) in a 50 μL system at 30°C for 14 hours. The reaction was terminated by heat inactivation at 65°C for 10 min. DNA purification was performed using phenol-chloroform-isoamyl alcohol (PCIA). Following debranching of the purification products, sequencing libraries were constructed using the NovaSeq X Series 25B Reagent kit according to the manufacturer’s protocol. Library quality was verified on an ABI QuantStudio 12K Flex, and final libraries were sequenced on an Illumina NovaSeq X Plus platform with 150-bp paired-end reads.

#### Quality control and raw data pre-processing

Both low quality and potential adapter sequences were filtered using Fastp^47^ (v0.12.4). Quality-controlled reads were subsequently aligned to the human reference genome (hg38), employing Burrows-Wheeler Aligner MEM (v0.7.17) for next-generation DNA sequencing data and minimap2^48^ (v2.1) for third-generation DNA sequencing data, both using default parameters. The resulting aligned BAM files were then utilized for microbial detection.

#### CIRCMIP: Circular DNA-based microbial detection and noise reduction

CIRCMIP (CIRCular DNA-based Microbiome Identification Pipeline) represents a robust computational framework engineered for the high-fidelity identification of bacteriome-derived circular DNA from human tissue sequencing datasets. We developed and applied this pipeline to systematically analyze intratumoral bacteriomes across eight cancer types. The CIRCMIP pipeline encompasses six core analytical modules, comprising: (1) stringent host DNA filtration, (2) high-confidence bacterial species identification, (3) bacterial diversity profiling, (4) tumor associated bacterial characterization, (5) machine learning-based biomarker discovery, and (6) bacterial functional and metabolic profiling, as detailed in following sections.

##### Stringent host DNA filtration

CIRCMIP initiated the workflow by computationally removing human-derived sequences from the aligned BAM files to minimize host DNA contamination. The resulting microbial-enriched reads were subsequently classified against kraken2DB (including Archaea, Bacteria, Viruses, Fungi, Human, and common laboratory contaminants) using Kraken2^49^ (v2.1.3) with the k-mer counting features of KrakenUniq for taxonomic assignment.

##### High-confidence bacterial species identification

Filter 1: General contaminant removal and normalization

Within the CIRCMIP framework, non-bacterial lineages (Archaea, Viruses, Fungi and human) and potential laboratory contaminants identified via kraken2DB were sequentially excluded from the KrakenUniq classification results (Filter 1). The remaining bacterial reads were normalized in two stages via total count scaling and conversion to reads per million bacterial mapped reads (RPBM) for robust inter-sample comparison.

Filter 2: Sample-level signal denoising

CIRCMIP further implemented a tripartite Spearman’s rank correlation framework (Filter 2) for secondary taxonomic refinement. This strategy was predicated on the robust positive associations characterizing genuine microbial signatures^50^. It evaluated three pairwise correlations: (1) total versus unique k-mer counts, (2) total k-mer counts versus total read counts, and (3) total read counts versus unique k-mer counts. Bacterial taxa fulfilling all three criteria (*P* < 0.05) were retained for subsequent analysis. In small cohorts (n ≤5), this filter was bypassed to account for limited statistical power.

Filter 3: Cell line-based contamination screening

CIRCMIP implemented a contamination-screening framework leveraging 2,491 sterile cell experiments as a reference benchmark^50^. For each candidate taxon, the relative abundance in test samples was compared against the 95th-percentile threshold derived from these negative controls. Taxa demonstrating significant enrichment beyond this baseline (one-sample quantile test, *P* < 0.05) were systematically excluded as putative laboratory contaminants. This cell line-anchored approach ensured the rigorous removal of non-biological noise while preserving authentic microbial signatures.

Filter 4: Healthy tissue-based taxonomic filtering

CIRCMIP utilized bacterial abundance profiles from 18 healthy controls, sequenced via identical circular DNA enrichment methods^51^, to establish method-specific baseline thresholds. This filter retained only those bacterial taxa with abundances in test samples exceeding the minimum observed levels in the healthy reference set. This strategy was designed to filter out pervasive microbial signatures inherent to the normal tissue environment and enrichment process.

Filter 5: Low-abundance taxonomic filtering

CIRCMIP applied a stringent abundance cutoff to minimize sequencing noise and stochastic artifacts. Bacterial taxa with a relative abundance below 0.01% were systematically excluded from the dataset. This threshold eliminated low-prevalence signatures likely representing technical noise rather than functional bacterial taxa, thereby preserving the core bacterial community structure for downstream analysis.

##### Bacterial diversity profiling

CIRCMIP evaluated bacterial alpha diversity (Shannon index) and beta diversity at both the genus and species levels utilizing the RPBM matrix. Alpha diversity was calculated with the diversity function, while beta diversity was computed using the vegdist function from the vegan R package (v2.6-4), utilizing both Bray-Curtis (quantitative) and Jaccard (qualitative) dissimilarities as distance metrics. To evaluate differences in beta diversity between tumor and NAT samples within each cancer type, we performed permutational multivariate analysis of variance (PERMANOVA) using the adonis2 function with 1,000 permutations. These variations were further visualized through Principal Coordinates Analysis (PCoA), and *P* < 0.05 was considered statistically significant. All analyses were conducted independently for bacterial compositions at both the genus and species levels. Pairwise cancer type similarities were further assessed by converting median dissimilarity scores to similarities, calculated as 1 minus Bray-Curtis for abundance-based profiles and 1 minus Jaccard for presence-absence patterns, followed by the comparison of median values.

##### Tumor associated bacterial characterization

CIRCMIP identified tumor-associated bacterial signatures by conducting differential abundance analysis via MaAsLin2^52^ (v1.12.0). Linear mixed-effect models were applied to log-transformed bacterial abundance data, incorporating cancer type, patient gender, and age as fixed effects, with sample ID included as a random effect to account for intra-individual variability. These multivariate adjustments were implemented for cancer types with available clinical metadata to ensure the robustness of the identified associations. The model was configured to prioritize bacterial taxa demonstrating significant associations for subsequent validation.

To determine clinical relevance, the association between *S. lugdunensis* colonization and BLCA overall survival was evaluated (Table S4). Using the survival and survminer R packages (v0.4.9), patients were stratified by bacterial signatures for analysis via Kaplan Meier curves and Cox proportional hazards models, with significance established at a P value below 0.05.

##### Machine learning-based biomarker discovery

CIRCMIP implemented a rigorous machine learning framework for bacterial biomarker discovery. Feature selection was performed using recursive feature elimination (RFE) via the Caret package^53^ (v6.0.94), followed by model construction with gradient boosting machines (GBM) as previously described^54^. All models were trained on Z-score normalized bacterial RPBM matrices across six specific comparisons at both genus and species levels: (i) BLCA versus LUAD tumors, (ii) BLCA versus non-BLCA tumors, (iii) LUAD versus non-LUAD tumors, (iv) BLCA tumors versus matched NATs, (v) LUAD tumors versus matched NATs, and (vi) early-stage (T1/T2) versus late-stage (T3/T4) BLCA tumors. To ensure robust training, comparisons involving minority classes with fewer than 20 samples were excluded. The analytical workflow proceeded in five stages. First, each comparison dataset was stratified into training (70%) and testing (30%) subsets using a fixed random seed for reproducibility. Second, sample-wise Z-score normalization was applied to the training data, with identical parameters utilized for the test sets. Third, model optimization was achieved via 5-fold cross-validation on training data through a grid search of GBM hyperparameters, including interaction depth (1 to 10), number of trees (100 to 3000), learning rate (0.01 to 1), and the minimum observations per node (5 to 15). Fourth, performance was assessed on the holdout test sets by calculating the AUROC and AUPR using the PRROC package (v1.3.1). Finally, variable importance was quantified using the permutation method from the Caret package to identify the most discriminative bacterial features.

##### Bacterial functional and metabolic profiling

CIRCMIP predicted bacterial functional metagenomes using PICRUSt2^55^ (v2.5.2) based on the RPBM abundance matrix. Functional pathways were reconstructed to delineate the metabolic potential of the tumor resident bacterial community. Comparative analysis was subsequently conducted to identify divergent metabolic signatures between BLCA samples characterized by the enrichment or deficiency of *S. lugdunensis*.

#### WGS based benchmarking of bacterial detection sensitivity

CIRCMIP was compared against whole genome sequencing (WGS) to evaluate its relative enrichment efficiency and genomic coverage (**Table S2**). WGS was performed on a representative cohort of tumor specimens prioritized by tissue availability. WGS libraries were prepared from 0.5 μg of input genomic DNA. DNA was sheared to 300-500 bp fragments using a Covaris LE220 ultrasonicator. Following end repair, A-tailing, and adapter ligation (NovaSeq X Series 25B Reagent kit), library quality was assessed on an ABI Quant Studio 12K Flex. Sequencing was performed on an illumina NovaSeq X Plus platform with 150-bp paired-end reads.

To ensure direct comparability between platforms, WGS raw data were processed through a computational framework identical to CIRCMIP. This parallel analytical approach enabled a systematic comparison of bacterial signal enrichment and taxonomic representation between circular DNA and total genomic DNA fractions.

#### Transcriptomic data acquisition and bioinformatics analysis

Total RNA was isolated from frozen tissues using TRIzol reagent (Thermo Fisher Scientific), with quality control performed on an Agilent 2100 Bioanalyzer, requiring an RNA Integrity Number (RIN) exceeding 5.0 for library preparation. Sequencing libraries were constructed from 500 ng input RNA using the MGIEasy RNA Library Prep Kit (MGI) and sequenced on the DNBSEQ-T1/T5 platform (BGI) with 100 bp paired end reads.

Raw reads were quality-filtered using Fastp^47^ (v. 0.12.4) with adapter trimming. Sequence alignment was performed via HISAT2^56^ (v2.2.1), where human reads were mapped to the GRCh38 reference genome and mouse reads were aligned to the mm10 genome. Transcript quantification was executed using featureCounts^57^ (v2.0.1) and StringTie^58^ (v2.2.1). Differential expression analysis was conducted using DESeq2 (v1.14.1), with significance established as an FDR below 0.05 and a log2 fold change exceeding 1. Functional enrichment of differentially expressed genes was performed via Metascape^59^. Furthermore, Gene Set Enrichment Analysis (GSEA) was conducted to identify enriched biological pathways and processes across the entire transcriptome.

To characterize the immune landscape, the CIBERSORT deconvolution algorithm was applied to estimate the relative proportions of major immune cell types across 80 matched BLCA transcriptomes. Spearman rank correlation analysis was subsequently performed to assess the relationship between *S. lugdunensis* abundance and tumor infiltrating immune cell populations.

#### Quantitation of microbes and joint analysis with RNA-seq data

Gene set variation analysis (GSVA) was performed on StringTie-normalized gene expression profiles using hallmark pathway gene sets from the Molecular Signatures Database (MSigDB v2023.2) to quantify tumor pathway activities. Concurrently, immune cell composition was deconvoluted from FPKM-normalized RNA-seq data using CIBERSORT^60^ (absolute mode) implemented through the immunedeconv R package (v2.0.3) with LM22 signature matrix. To investigate microbiome-immune-pathway relationships, we employed Adaptive Microbiome α-diversity Association Analysis (aMiAD v2.0)^61^ with Gaussian kernel, where rarefied α-diversity indices (Shannon diversity, species richness, and Gini-Simpson index) served as response variables, and cancer type/gender/age-adjusted residuals of pathway activities (from GSVA) were included as predictors, with statistical significance assessed through 5,000 permutations to control for multiple testing.

#### Spatial validation via RNAscope and immunofluorescence

To visualize the spatial distribution of bacteria within the tumor architecture, we performed a combined protocol involving RNAscope in situ hybridization and immunofluorescence on formalin fixed paraffin embedded tissue sections. Bacterial 16S rRNA was detected using the RNAscope Multiplex Fluorescent Reagent Kit v2 (Advanced Cell Diagnostics, Cat# 323100) in conjunction with the RNA Protein Co Detection Ancillary Kit (Cat# 323180) to ensure the integrity of both nucleic acid and protein signals. Specifically, tissue sections were hybridized with the RNAscope Probe EB 16S rRNA (Cat# 464461), a universal bacterial probe. Following the RNA hybridization procedure, tumor epithelial cells were identified via immunostaining using a PanCK primary antibody (Abcam, Cat# ab7753) followed by incubation with an Alexa Fluor 488 conjugated secondary antibody. Nuclei were counterstained utilizing DAPI at a concentration of 10 micrograms per milliliter (Solarbio, Cat# C0065). High resolution images were captured using a Zeiss LSM 900 laser scanning confocal microscope utilizing a 63 times oil immersion objective.

#### Bacterial metabolite preparation and targeted lipidomic profiling

*S. lugdunensis* was cultured in BHI broth at 37°C for 48 hours under aerobic conditions. After that, extract the culture supernatant with an equal volume of ethyl acetate (1:1 ratio). The upper organic phase was collected and evaporated to dryness to obtain the crude extract. This crude extract was then dissolved in DMSO (Sigma-Aldrich, Cat# D2650; CAS: 67-68-5) for subsequent cell treatments. Parallel BHI medium processing without bacterial inoculation served as the negative control.

For tissue processing, frozen clinical specimens consisting of *S. lugdunensis* positive and negative tumors along with matched normal adjacent tissues were weighed and homogenized in chilled buffer under ice cold conditions to generate standardized tissue homogenates. These homogenates, together with the bacterial lysates and BHI controls, were subsequently subjected to a deep targeted lipidomics platform for effector molecule screening.

Mass spectrometry analysis was conducted via ultrahigh performance liquid chromatography tandem mass spectrometry. The targeted lipidomics panel was specifically designed to detect and quantify various lipid species across all sample types (**Table S5**). Subsequent bioinformatic evaluations were executed on normalized datasets using MetaboAnalyst^62^ (v.5).

#### Real-time cell proliferation assay

T24 cells (2,000 cells/well) were seeded in 96-well plates and exposed to live *S. lugdunensis* (BNCC, Cat# BNCC262227) at a multiplicity of infection (MOI) of 100. For lipid stimulations, cells were incubated with a concentration gradient (0-45 µM, 5 µM increments) of LPC14:0 (Cat# 855575; CAS: 20559-16-4), LPC16:0 (Cat# 855675; CAS: 17364-16-8), EggLPC (Cat# 830071; CAS: 97281-36-2), or PC14:0 (Cat# 850345; CAS: 18194-24-6, all from Avanti Polar Lipids). Brain Heart Infusion (BHI) medium and respective lipid vehicles served as parallel controls.

Cell proliferation was continuously monitored for 69-75 h using the Incucyte S3 Live-Cell Analysis System (Sartorius). Phase-contrast images were acquired at 3h intervals. The percent confluence was quantified using the Incucyte Base Analysis Software (version 2024, Sartorius) and normalized to baseline measurements obtained at 0 h (Table S5). Additionally, RNA-seq was performed on T24 cells following LPC 14:0 treatment using the DNBSEQ-T7 platform (BGI) with 150-bp paired-end sequencing. The raw data underwent quality control, mapping, and differential analysis consistent with the aforementioned transcriptomic pipelines.

#### Bacterial challenge and metabolite administration

For the bacterial intervention study, mice were randomly assigned to three groups: (1) vehicle control, (2) non-pathogenic *Escherichia coli* (*E. col*; this study, laboratory isolate), and (3) *S. lugdunensis* (BNCC, Cat# BNCC262227). Following the initial tumor cell injection, bacterial strains were administered via intragastric gavage at a dosage of 1 × 10^5^ colony-forming units (CFU) per mouse every 5 days. To ensure local colonization, an intratumoral injection of the corresponding bacteria or vehicle was performed on day 5.

For the lipid metabolite intervention, mice were randomly allocated into three groups: vehicle control, LPC14:0, and LPC16:0. The lipid metabolites were administered via intratumoral injection (100 µL per mouse) every 2 days, starting from day 5. Tumor dimensions (length and width) were measured using digital calipers every 5 days.

Tumor volume was calculated using the standard formula 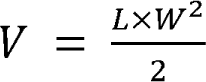, where *V* represents tumor volume, *L* represents tumor length, and *W* represents tumor width. Mouse body weights were monitored longitudinally throughout the duration of the study.

At the experimental endpoint, tumor tissues from the *S. lugdunensis*-infected, LPC 14:0 and LPC16:0-treated groups were harvested for transcriptomic profiling. Total RNA was extracted and sequenced on the DNBSEQ-T7 platform (BGI) using 150-bp paired-end reads. Bioinformatics analysis, including data filtering, alignment, and identification of differentially expressed genes, was conducted using the same pipeline as established for the *in vitro* RNA-seq analysis.

#### Intracellular Bacterial Infection Assay

The experimental steps were performed as previously reported (PMID: 39846310). The *S. lugdunensis* strain was stained with HADA hydrochloride (MedChemExpress, Cat# HY-131045, CAS No.: 2253733-10-5) for 30 minutes. The fluorescently labeled bacterial strain was then added at a 1:2 ratio to infect T24 cells for 2 hours. The fluorescently labeled bacterial strain was then added at a 1:2 ratio to infect T24 cells for 2 hours, followed by immunofluorescence staining and imaging. Brifly, the cells were fixed with 4% paraformaldehyde, permeabilized with 0.5% Triton X-100, and blocked with 1% BSA. They were then stained with the Actin-Tracker Red-594 (Beyotime, Cat# C2205S) and imaged using a confocal microscope.

#### Tissue processing and immunohistochemistry

At day 30, mice were euthanized and tumors were harvested, weighed, and photographed. For histological analysis, tumor tissues were fixed in four percent paraformaldehyde, embedded in paraffin, and sectioned at a thickness of five microns. Immunohistochemistry was performed to evaluate cellular proliferation using a Ki67 primary antibody (HUABIO, Cat# HA721115). Sections were counterstained with hematoxylin and visualized under a light microscope. The percentage of Ki67 positive cells was quantified by counting positive nuclei in five randomly selected high power fields per section using ImageJ software.

#### Fatty acid uptake assay

T24 bladder cancer cells (1×10^6^ cells/well) were treated with 30 µM LPC14:0 or an ethanol vehicle for 18 hours. Fatty acid uptake capacity was then quantified using a fluorometric assay kit (Elabscience, Cat# E-BC-F067). Briefly, cells were incubated with a fluorescent fatty acid analog at 37°C for 30 min, while parallel background controls were maintained in substrate-free buffer. Post-incubation, cells were washed, harvested by trypsinization, and resuspended for fluorescence measurement (Ex/Em 485/515 nm). To ensure rigorous quantification, total protein concentration was determined via a BCA assay (Applygen, Cat# P1511) to account for potential variations in cell density. Relative fatty acid uptake was calculated as background-subtracted fluorescence intensity (ΔRFU) normalized to the total protein content of each respective sample.

#### Fatty acid oxidation (FAO) assay

FAO capacity was assessed using a Fatty Acid Oxidation (FAO) Colorimetric Assay Kit (Elabscience, Cat# E-BC-K784-M) in T24 cells (treated similarly to those in the fatty acid uptake assay) and in subcutaneous tumors harvested from the *in vivo* models. For the tumor tissues, samples were collected from mice receiving either bacterial treatments (non-pathogenic *E. coli*, *S. lugdunensis*, or vehicle control) or lipid metabolite administrations (LPC 14:0, LPC 16:0, or vehicle control).

Samples were homogenized in an extraction buffer and centrifuged at 14,000 × g for 10 min to obtain the supernatants. These extracts were incubated at 37°C for 30 min with chromogenic and substrate reagents, while parallel background controls received a buffer instead of the substrate to account for endogenous background signals. The net absorbance change at 450 nm (ΔA_450_) was monitored against an NADH standard curve. Final FAO activity (U/g protein) was normalized to the total protein concentration of each sample, as determined using a BCA Protein Assay Kit (Applygen, Cat# P1511).

#### LPC14:0 probe design and synthesis

A bifunctional photo-crosslinking probe, designated as *LPC14:0 (C_29_H_52_N_3_O_8_P, MW: 601.71 Da), was synthesized by esterifying the *sn*-2 hydroxyl group of native LPC 14:0 (C_22_H_46_NO_7_P, MW: 467.58 Da) with 2-(3-(but-3-yn-1-yl)-3H-diazirin-3-yl) acetic acid (MW: 152.15 Da). This customized probe incorporates a diazirine moiety for ultraviolet (UV)-induced covalent cross-linking to target interacting proteins and a terminal alkyne group for downstream bioorthogonal enrichment via click chemistry.

#### In situ photo-crosslinking and bioorthogonal capture

T24 cells (QuiCell, Cat# QuiCell-T395) were incubated with the LPC14:0 probe at a final concentration of 30 µM for 18 h at 37°C to allow for in situ target engagement. Following incubation, cells were washed with ice-cold PBS and irradiated with UV light (365 nm) for 15 min on ice to induce the covalent cross-linking of interacting proteins. Cells were subsequently harvested and lysed in 300 µL of enhanced RIPA lysis buffer (Applygen, Cat# C1053+) supplemented with protease inhibitors. For downstream target enrichment, the cross-linked lysates were subjected to copper-catalyzed azide-alkyne cycloaddition (CuAAC) to conjugate the probe with a biotin-azide tag. This affinity enrichment was performed utilizing the ChomiX®-PulldownKit–Streptavidin Magnetic Beads-Based Protein Enrichment Kit (Chomix, Cat# 02030011) according to the manufacturer’s instructions.

#### In situ fluorescence visualization and competitive binding

To visualize the specific engagement of the lipid probe with its intracellular targets, in situ fluorescence labeling and competitive binding assays were performed. T24 cells were seeded in 6-well plates and randomly assigned to nine experimental groups: (1) blank control, (2) 30 µM native LPC14:0, (3) 10 µM LPC14:0 probe and (4) 10 µM LPC14:0 probe co-incubated with a 10-fold excess (100 µM) of native LPC14:0 for competitive target validation.

Following the designated treatments, cells were washed with ice-cold PBS and irradiated with UV light (365 nm) for 15 min on ice to induce the covalent photo-crosslinking of interacting proteins. Cells were subsequently fixed with 4% paraformaldehyde (Solarbio, Cat# P1110) at room temperature and permeabilized. For fluorescence visualization, copper-catalyzed azide-alkyne cycloaddition (CuAAC) was performed utilizing the BeyoClick™ EdU-594 Cell Proliferation Assay Kit (Beyotime), with the EdU incorporation step strictly omitted. The cross-linked lipid probes were conjugated with the provided Azide 594. High-resolution fluorescence images were ultimately acquired utilizing a ZEISS LSM 980 confocal laser scanning microscope..

#### Affinity purification and target identification via LC-MS/MS

To identify specific interacting protein targets, T24 cells were incubated with 30 µM of the LPC14:0 probe or an equivalent volume of the vehicle control for 18 h. Following *in situ* photo-crosslinking and cell lysis, the proteome was subjected to CuAAC with a biotin-azide probe. Biotinylated proteins were enriched using streptavidin-coated magnetic beads.

After extensive washing to remove non-specific binders, the enriched protein complexes were eluted from the beads by boiling in standard SDS loading buffer. The eluents were then loaded onto precast polyacrylamide gels (GenScript, Cat# M01158) and resolved by SDS-PAGE at a constant voltage of 80 V for 30 min. For downstream mass spectrometry analysis, the entire protein lane corresponding to each sample was excised from the gel. The excised gel bands were subjected to standard in-gel tryptic digestion. The resulting peptide mixtures were extracted, desalted, and analyzed by high-resolution liquid chromatography-tandem mass spectrometry (LC-MS/MS). Specific protein targets were subsequently identified by comparing the mass spectrometry enrichment profiles of the probe-treated group against the vehicle control, effectively filtering out non-specific background signals.

#### Molecular docking simulation of the LPC14:0-PPAR**δ** binding interface

The binding mode between *S. lugdunensis*-derived LPC14:0 and human PPARδ was characterized using molecular docking simulations. The high-resolution crystal structure of the human PPARδ ligand-binding domain (LBD) in complex with eicosapentaenoic acid (EPA) was retrieved from the Protein Data Bank (PDB ID: 1GWX). Preparation of the receptor was performed using PyMOL (Schrödinger, LLC) to systematically remove the co-crystallized ligand, water molecules, and ions. For ligand preparation, the chemical structure of LPC14:0 was obtained from the ZINC database and subjected to energy minimization using ChemDraw 3D (PerkinElmer) to achieve a thermodynamically stable three-dimensional. Blind docking was executed using the CB-DOCK2 server to predict the most favorable binding pockets and orientations. The resulting docking poses were evaluated based on their binding affinity scores and the consistency of the interactions with critical residues within the PPARδ ligand-binding pocket.

### QUANTIFICATION AND STATISTICAL ANALYSIS

Data are presented as the mean ± standard error of the mean (SEM). Asterisks in the figures indicate the level of statistical significance (SEM). Asterisks in the figures indicate the level of statistical significance (* *P* < 0.05, ** *P* < 0.01, *** *P* < 0.001) as determined using either a two-tailed unpaired Student’s t test or a Mann-Whitney test, as explicitly defined in the respective figure captions. All statistical evaluations and data visualizations were conducted utilizing R software (version 4.5.2), with a p value below 0.05 considered to be statistically significant.

For in situ fluorescence visualization, image processing was conducted utilizing ImageJ/Fiji software (NIH). To ensure rigorous visual comparability across different experimental conditions, the fluorescence intensity display ranges (brightness and contrast) were uniformly standardized for all captured images. This standardized adjustment was applied consistently across all groups to faithfully represent the specific probe signals and minimize subjective bias during visual comparison. The processed representative images were subsequently assembled for qualitative presentation.

## REFERENCES

1. de Martel, C., Ferlay, J., Franceschi, S., Vignat, J., Bray, F., Forman, D., and Plummer, M. (2012). Global burden of cancers attributable to infections in 2008: a review and synthetic analysis. Lancet Oncol 13, 607–615. 10.1016/S1470-2045(12)70137-7.

2. Wong, S.H., Zhao, L., Zhang, X., Nakatsu, G., Han, J., Xu, W., Xiao, X., Kwong, T.N.Y., Tsoi, H., Wu, W.K.K., et al. (2017). Gavage of Fecal Samples From Patients With Colorectal Cancer Promotes Intestinal Carcinogenesis in Germ-Free and Conventional Mice. Gastroenterology 153, 1621–1633 e1626. 10.1053/j.gastro.2017.08.022.

3. Kostic, A.D., Chun, E., Robertson, L., Glickman, J.N., Gallini, C.A., Michaud, M., Clancy, T.E., Chung, D.C., Lochhead, P., Hold, G.L., et al. (2013). Fusobacterium nucleatum potentiates intestinal tumorigenesis and modulates the tumor-immune microenvironment. Cell Host Microbe 14, 207–215. 10.1016/j.chom.2013.07.007.

4. Gopalakrishnan, V., Spencer, C.N., Nezi, L., Reuben, A., Andrews, M.C., Karpinets, T.V., Prieto, P.A., Vicente, D., Hoffman, K., Wei, S.C., et al. (2018). Gut microbiome modulates response to anti-PD-1 immunotherapy in melanoma patients. Science 359, 97–103. 10.1126/science.aan4236.

5. Matson, V., Fessler, J., Bao, R., Chongsuwat, T., Zha, Y., Alegre, M.L., Luke, J.J., and Gajewski, T.F. (2018). The commensal microbiome is associated with anti-PD-1 efficacy in metastatic melanoma patients. Science 359, 104–108. 10.1126/science.aao3290.

6. Routy, B., Le Chatelier, E., Derosa, L., Duong, C.P.M., Alou, M.T., Daillere, R., Fluckiger, A., Messaoudene, M., Rauber, C., Roberti, M.P., et al. (2018). Gut microbiome influences efficacy of PD-1-based immunotherapy against epithelial tumors. Science 359, 91–97. 10.1126/science.aan3706.

7. Viaud, S., Saccheri, F., Mignot, G., Yamazaki, T., Daillere, R., Hannani, D., Enot, D.P., Pfirschke, C., Engblom, C., Pittet, M.J., et al. (2013). The intestinal microbiota modulates the anticancer immune effects of cyclophosphamide. Science 342, 971–976. 10.1126/science.1240537.

8. Iida, N., Dzutsev, A., Stewart, C.A., Smith, L., Bouladoux, N., Weingarten, R.A., Molina, D.A., Salcedo, R., Back, T., Cramer, S., et al. (2013). Commensal bacteria control cancer response to therapy by modulating the tumor microenvironment. Science 342, 967–970. 10.1126/science.1240527.

9. Helmink, B.A., Khan, M.A.W., Hermann, A., Gopalakrishnan, V., and Wargo, J.A. (2019). The microbiome, cancer, and cancer therapy. Nat Med 25, 377–388. 10.1038/s41591-019-0377-7.

10. Jiang, J., Mei, J., Ma, Y., Jiang, S., Zhang, J., Yi, S., Feng, C., Liu, Y., and Liu, Y. (2022). Tumor hijacks macrophages and microbiota through extracellular vesicles. Exploration (Beijing) 2, 20210144. 10.1002/EXP.20210144.

11. Su, J., Song, Y., Zhu, Z., Huang, X., Fan, J., Qiao, J., and Mao, F. (2024). Cell-cell communication: new insights and clinical implications. Signal Transduct Target Ther 9, 196. 10.1038/s41392-024-01888-z.

12. Wang, T., Zhao, H., Sun, X., Ding, Y., Zhu, Z., Yu, X., Zhu, R., Wang, D., Li, K., Liu, Y., et al. (2026). Pancancer Fine-Mapping of Mutational Intolerance Identifies CHEK1 as an Immunosuppressive Driver in Lung Adenocarcinoma. Adv Sci (Weinh), e21265. 10.1002/advs.202521265.

13. Zhu, Z., Zhang, M., Song, Y., Jiang, W., Cao, F., Yao, Y., Yu, X., Zhao, H., Baiyin, H., Chang, D., et al. (2026). Granulin+ Macrophages Promote Lineage Plasticity in Prostate Cancer Through Paracrine Signaling Loops. Genomics Proteomics Bioinformatics. 10.1093/gpbjnl/qzag024.

14. Xu, X., Su, J., Zhu, R., Li, K., Zhao, X., Fan, J., and Mao, F. (2025). From morphology to single-cell molecules: high-resolution 3D histology in biomedicine. Mol Cancer 24, 63. 10.1186/s12943-025-02240-x.

15. Nejman, D., Livyatan, I., Fuks, G., Gavert, N., Zwang, Y., Geller, L.T., Rotter-Maskowitz, A., Weiser, R., Mallel, G., Gigi, E., et al. (2020). The human tumor microbiome is composed of tumor type-specific intracellular bacteria. Science 368, 973–980. 10.1126/science.aay9189.

16. Ma, J., Huang, L., Hu, D., Zeng, S., Han, Y., and Shen, H. (2021). The role of the tumor microbe microenvironment in the tumor immune microenvironment: bystander, activator, or inhibitor? J Exp Clin Cancer Res 40, 327. 10.1186/s13046-021-02128-w.

17. Niu, Y., Feng, J., Ma, J., Xiao, T., and Yuan, W. (2025). Targeting the Tumor Microbiota in Cancer Therapy Basing on Nanomaterials. Exploration (Beijing) 5, e20210185. 10.1002/EXP.20210185.

18. Davis, N.M., Proctor, D.M., Holmes, S.P., Relman, D.A., and Callahan, B.J. (2018). Simple statistical identification and removal of contaminant sequences in marker-gene and metagenomics data. Microbiome 6, 226. 10.1186/s40168-018-0605-2.

19. Eisenhofer, R., Minich, J.J., Marotz, C., Cooper, A., Knight, R., and Weyrich, L.S. (2019). Contamination in Low Microbial Biomass Microbiome Studies: Issues and Recommendations. Trends Microbiol 27, 105–117. 10.1016/j.tim.2018.11.003.

20. Glassing, A., Dowd, S.E., Galandiuk, S., Davis, B., and Chiodini, R.J. (2016). Inherent bacterial DNA contamination of extraction and sequencing reagents may affect interpretation of microbiota in low bacterial biomass samples. Gut Pathog 8, 24. 10.1186/s13099-016-0103-7.

21. Kostic, A.D., Gevers, D., Pedamallu, C.S., Michaud, M., Duke, F., Earl, A.M., Ojesina, A.I., Jung, J., Bass, A.J., Tabernero, J., et al. (2012). Genomic analysis identifies association of Fusobacterium with colorectal carcinoma. Genome Res 22, 292–298. 10.1101/gr.126573.111.

22. Bullman, S., Pedamallu, C.S., Sicinska, E., Clancy, T.E., Zhang, X., Cai, D., Neuberg, D., Huang, K., Guevara, F., Nelson, T., et al. (2017). Analysis of Fusobacterium persistence and antibiotic response in colorectal cancer. Science 358, 1443–1448. 10.1126/science.aal5240.

23. Flanagan, L., Schmid, J., Ebert, M., Soucek, P., Kunicka, T., Liska, V., Bruha, J., Neary, P., Dezeeuw, N., Tommasino, M., et al. (2014). Fusobacterium nucleatum associates with stages of colorectal neoplasia development, colorectal cancer and disease outcome. Eur J Clin Microbiol Infect Dis 33, 1381–1390. 10.1007/s10096-014-2081-3.

24. Yu, T., Guo, F., Yu, Y., Sun, T., Ma, D., Han, J., Qian, Y., Kryczek, I., Sun, D., Nagarsheth, N., et al. (2017). Fusobacterium nucleatum Promotes Chemoresistance to Colorectal Cancer by Modulating Autophagy. Cell 170, 548–563 e516. 10.1016/j.cell.2017.07.008.

25. Robinson, K.M., Crabtree, J., Mattick, J.S., Anderson, K.E., and Dunning Hotopp, J.C. (2017). Distinguishing potential bacteria-tumor associations from contamination in a secondary data analysis of public cancer genome sequence data. Microbiome 5, 9. 10.1186/s40168-016-0224-8.

26. Yao, B., Liu, X., Ruan, K., Fang, X., Jiang, C., Bian, W., Guo, Y., Zhu, X., Shang, Z., Hu, T., et al. (2026). Divergent tumor immunity determined by bacteria-cancer cell engagement. Cell 189, 1748–1767 e1726. 10.1016/j.cell.2025.12.044.

27. Yu, H., Du, Y., He, Y., Sun, Y., Li, J., Jia, B., Chen, J., Peng, X., An, T., Li, J., et al. (2025). Lactate production by tumor-resident Staphylococcus promotes metastatic colonization in lung adenocarcinoma. Cell Host Microbe 33, 1089–1105 e1087. 10.1016/j.chom.2025.06.013.

28. Chen, C., Gao, Y., Lu, R., Wu, Y., Yao, Y., Sun, R., Wu, J., Ji, P., Quan, W., Wu, D., et al. (2025). TERC Stimulates Fatty Acid Metabolism to Promote Bladder Cancer Progression. Cancer Res 85, 3689–3705. 10.1158/0008-5472.CAN-24-3439.

29. Cai, J., Shi, J., Chen, C., He, M., Wang, Z., and Liu, Y. (2023). Structural-Activity Relationship-Inspired the Discovery of Saturated Fatty Acids as Novel Colistin Enhancers. Adv Sci (Weinh) 10, e2302182. 10.1002/advs.202302182.

30. den Hartog, G., Butcher, L.D., Ablack, A.L., Pace, L.A., Ablack, J.N.G., Xiong, R., Das, S., Stappenbeck, T.S., Eckmann, L., Ernst, P.B., and Crowe, S.E. (2020). Apurinic/Apyrimidinic Endonuclease 1 Restricts the Internalization of Bacteria Into Human Intestinal Epithelial Cells Through the Inhibition of Rac1. Front Immunol 11, 553994. 10.3389/fimmu.2020.553994.

31. Thuenauer, R., Kuhn, K., Guo, Y., Kotsis, F., Xu, M., Trefzer, A., Altmann, S., Wehrum, S., Heshmatpour, N., Faust, B., et al. (2022). The Lectin LecB Induces Patches with Basolateral Characteristics at the Apical Membrane to Promote Pseudomonas aeruginosa Host Cell Invasion. mBio 13, e0081922. 10.1128/mbio.00819-22.

32. Tajima, N., Simorowski, N., Yovanno, R.A., Regan, M.C., Michalski, K., Gomez, R., Lau, A.Y., and Furukawa, H. (2022). Development and characterization of functional antibodies targeting NMDA receptors. Nat Commun 13, 923. 10.1038/s41467-022-28559-3.

33. Battaglia, T.W., Mimpen, I.L., Traets, J.J.H., van Hoeck, A., Zeverijn, L.J., Geurts, B.S., de Wit, G.F., Noe, M., Hofland, I., Vos, J.L., et al. (2024). A pan-cancer analysis of the microbiome in metastatic cancer. Cell 187, 2324–2335 e2319. 10.1016/j.cell.2024.03.021.

34. Banerjee, S., Tian, T., Wei, Z., Shih, N., Feldman, M.D., Alwine, J.C., Coukos, G., and Robertson, E.S. (2017). The ovarian cancer oncobiome. Oncotarget 8, 36225–36245. 10.18632/oncotarget.16717.

35. Greathouse, K.L., White, J.R., Vargas, A.J., Bliskovsky, V.V., Beck, J.A., von Muhlinen, N., Polley, E.C., Bowman, E.D., Khan, M.A., Robles, A.I., et al. (2018). Interaction between the microbiome and TP53 in human lung cancer. Genome Biol 19, 123. 10.1186/s13059-018-1501-6.

36. Yu, G., Gail, M.H., Consonni, D., Carugno, M., Humphrys, M., Pesatori, A.C., Caporaso, N.E., Goedert, J.J., Ravel, J., and Landi, M.T. (2016). Characterizing human lung tissue microbiota and its relationship to epidemiological and clinical features. Genome Biol 17, 163. 10.1186/s13059-016-1021-1.

37. Fu, A., Yao, B., Dong, T., Chen, Y., Yao, J., Liu, Y., Li, H., Bai, H., Liu, X., Zhang, Y., et al. (2022). Tumor-resident intracellular microbiota promotes metastatic colonization in breast cancer. Cell 185, 1356–1372 e1326. 10.1016/j.cell.2022.02.027.

38. Thomas, V.M., Brown, R.M., Ashcraft, D.S., and Pankey, G.A. (2019). Synergistic effect between nisin and polymyxin B against pandrug-resistant and extensively drug-resistant Acinetobacter baumannii. Int J Antimicrob Agents 53, 663–668. 10.1016/j.ijantimicag.2019.03.009.

39. Hu, D., Chen, W., Wang, W., Tian, D., Fu, P., Ren, P., Mu, Q., Li, G., and Jiang, X. (2023). Hypercapsule is the cornerstone of Klebsiella pneumoniae in inducing pyogenic liver abscess. Front Cell Infect Microbiol 13, 1147855. 10.3389/fcimb.2023.1147855.

40. Debelius, J.W., Huang, T., Cai, Y., Ploner, A., Barrett, D., Zhou, X., Xiao, X., Li, Y., Liao, J., Zheng, Y., et al. (2020). Subspecies Niche Specialization in the Oral Microbiome Is Associated with Nasopharyngeal Carcinoma Risk. mSystems 5. 10.1128/mSystems.00065-20.

41. Li, C., Hu, Z., Zhang, W., Ji, Y., Wang, F., Zeng, Y., Sinha, S., Mao, F., Lin, C., and Lv, W. (2026). Extrachromosomal DNA in urothelial carcinoma: mechanisms and clinical applications. Nat Rev Urol. 10.1038/s41585-026-01134-x.

42. Lv, W., Pan, X., Han, P., Wu, S., Zeng, Y., Wang, Q., Guo, L., Xu, M., Qi, Y., Deng, L., et al. (2024). Extrachromosomal circular DNA orchestrates genome heterogeneity in urothelial bladder carcinoma. Theranostics 14, 5102–5122. 10.7150/thno.99563.

43. Luo, X., Zhang, L., Cui, J., An, Q., Li, H., Zhang, Z., Sun, G., Huang, W., Li, Y., Li, C., et al. (2023). Small extrachromosomal circular DNAs as biomarkers for multi-cancer diagnosis and monitoring. Clin Transl Med 13, e1393. 10.1002/ctm2.1393.

44. Wu, N., Wei, L., Liu, Q., He, T., Huang, C., Jiang, Y., Li, K., Guo, H., Mao, F., and Zhao, X. (2025). Extrachromosomal circular DNA expressing miRNA promotes ovarian cancer progression. Clin Transl Med 15, e70445. 10.1002/ctm2.70445.

45. Jiang, X., Pan, X., Li, W., Han, P., Yu, J., Li, J., Zhang, H., Lv, W., Zhang, Y., He, Y., and Xiang, X. (2023). Genome-wide characterization of extrachromosomal circular DNA in gastric cancer and its potential role in carcinogenesis and cancer progression. Cell Mol Life Sci 80, 191. 10.1007/s00018-023-04838-0.

46. Zhu, Y., Liu, Z., Guo, Y., Li, S., Qu, Y., Dai, L., Chen, Y., Ning, W., Zhang, H., and Ma, L. (2022). Whole-genome sequencing of extrachromosomal circular DNA of cerebrospinal fluid of medulloblastoma. Front Oncol 12, 934159. 10.3389/fonc.2022.934159.

47. Chen, S., Zhou, Y., Chen, Y., and Gu, J. (2018). fastp: an ultra-fast all-in-one FASTQ preprocessor. Bioinformatics 34, i884–i890. 10.1093/bioinformatics/bty560.

48. Li, H. (2018). Minimap2: pairwise alignment for nucleotide sequences. Bioinformatics 34, 3094–3100. 10.1093/bioinformatics/bty191.

49. Wood, D.E., Lu, J., and Langmead, B. (2019). Improved metagenomic analysis with Kraken 2. Genome Biol 20, 257. 10.1186/s13059-019-1891-0.

50. Ghaddar, B., Biswas, A., Harris, C., Omary, M.B., Carpizo, D.R., Blaser, M.J., and De, S. (2022). Tumor microbiome links cellular programs and immunity in pancreatic cancer. Cancer Cell 40, 1240–1253 e1245. 10.1016/j.ccell.2022.09.009.

51. Mann, L., Seibt, K.M., Weber, B., and Heitkam, T. (2022). ECCsplorer: a pipeline to detect extrachromosomal circular DNA (eccDNA) from next-generation sequencing data. BMC Bioinformatics 23, 40. 10.1186/s12859-021-04545-2.

52. Mallick, H., Rahnavard, A., McIver, L.J., Ma, S., Zhang, Y., Nguyen, L.H., Tickle, T.L., Weingart, G., Ren, B., Schwager, E.H., et al. (2021). Multivariable association discovery in population-scale meta-omics studies. PLoS Comput Biol 17, e1009442. 10.1371/journal.pcbi.1009442.

53. Kuhn, M. (2008). Building predictive models in R using the caret package. J. Stat. Softw 28.

54. Wang, H., Wang, T., Zhao, X., Wu, H., You, M., Sun, Z., and Mao, F. (2020). AI-Driver: an ensemble method for identifying driver mutations in personal cancer genomes. NAR Genom Bioinform 2, lqaa084. 10.1093/nargab/lqaa084.

55. Douglas, G.M., Maffei, V.J., Zaneveld, J.R., Yurgel, S.N., Brown, J.R., Taylor, C.M., Huttenhower, C., and Langille, M.G.I. (2020). PICRUSt2 for prediction of metagenome functions. Nat Biotechnol 38, 685–688. 10.1038/s41587-020-0548-6.

56. Kim, D., Paggi, J.M., Park, C., Bennett, C., and Salzberg, S.L. (2019). Graph-based genome alignment and genotyping with HISAT2 and HISAT-genotype. Nat Biotechnol 37, 907–915. 10.1038/s41587-019-0201-4.

57. Liao, Y., Smyth, G.K., and Shi, W. (2014). featureCounts: an efficient general purpose program for assigning sequence reads to genomic features. Bioinformatics 30, 923–930. 10.1093/bioinformatics/btt656.

58. Pertea, M., Pertea, G.M., Antonescu, C.M., Chang, T.C., Mendell, J.T., and Salzberg, S.L. (2015). StringTie enables improved reconstruction of a transcriptome from RNA-seq reads. Nat Biotechnol 33, 290–295. 10.1038/nbt.3122.

59. Zhou, Y., Zhou, B., Pache, L., Chang, M., Khodabakhshi, A.H., Tanaseichuk, O., Benner, C., and Chanda, S.K. (2019). Metascape provides a biologist-oriented resource for the analysis of systems-level datasets. Nat Commun 10, 1523. 10.1038/s41467-019-09234-6.

60. Sturm, G., Finotello, F., and List, M. (2020). Immunedeconv: An R Package for Unified Access to Computational Methods for Estimating Immune Cell Fractions from Bulk RNA-Sequencing Data. Methods Mol Biol 2120, 223–232. 10.1007/978-1-0716-0327-7_16.

61. Koh, H. (2018). An adaptive microbiome alpha-diversity-based association analysis method. Sci Rep 8, 18026. 10.1038/s41598-018-36355-7.

62. Xia, J., Psychogios, N., Young, N., and Wishart, D.S. (2009). MetaboAnalyst: a web server for metabolomic data analysis and interpretation. Nucleic Acids Res 37, W652–660. 10.1093/nar/gkp356.

