## Supplementary figures and images for "Pan-cancer circular genomics identifies intratumoral *Staphylococcus lugdunensis* as a metabolic driver in bladder cancer"

### Supplemental Figure 1

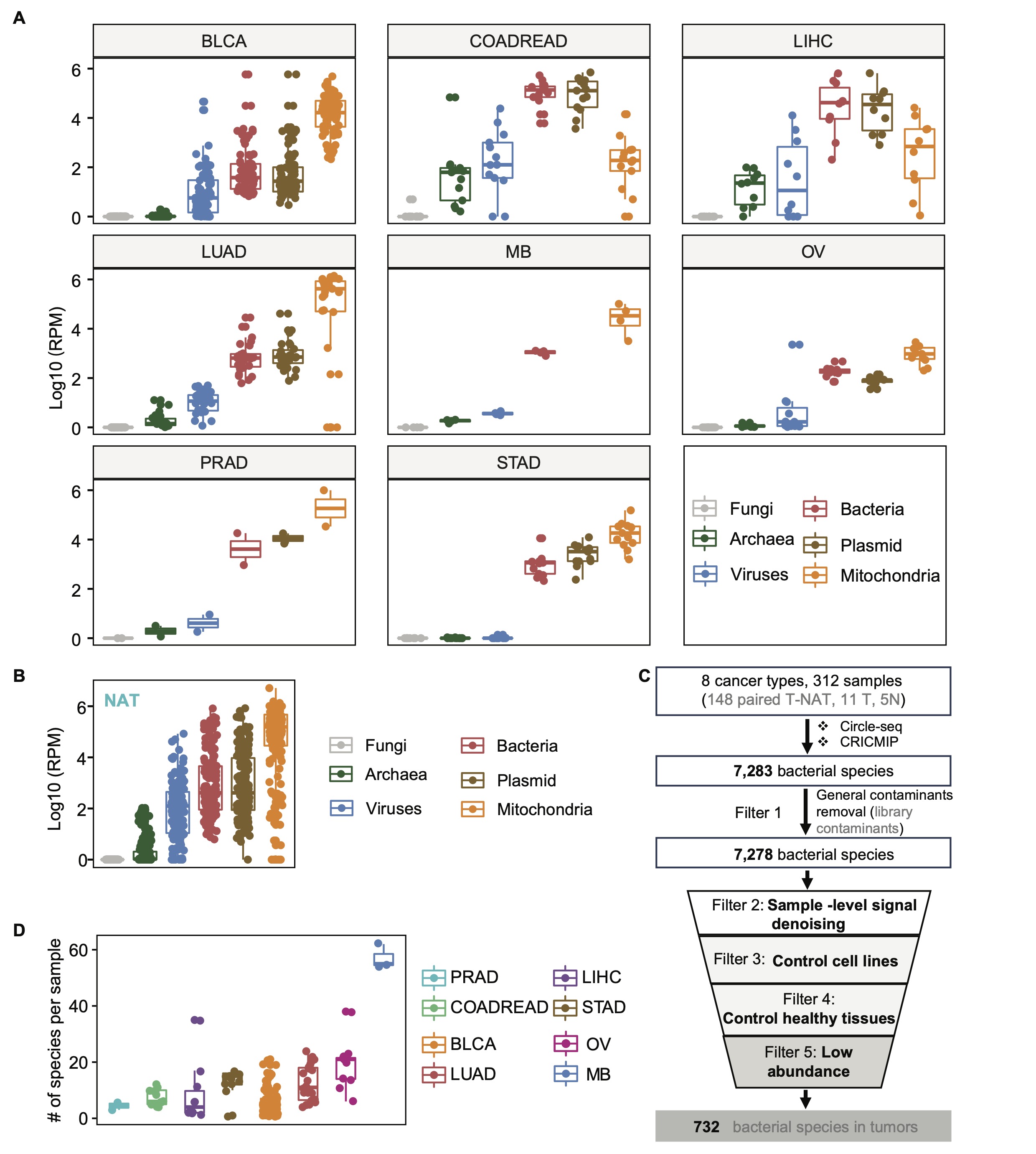

### Supplemental Figure 2

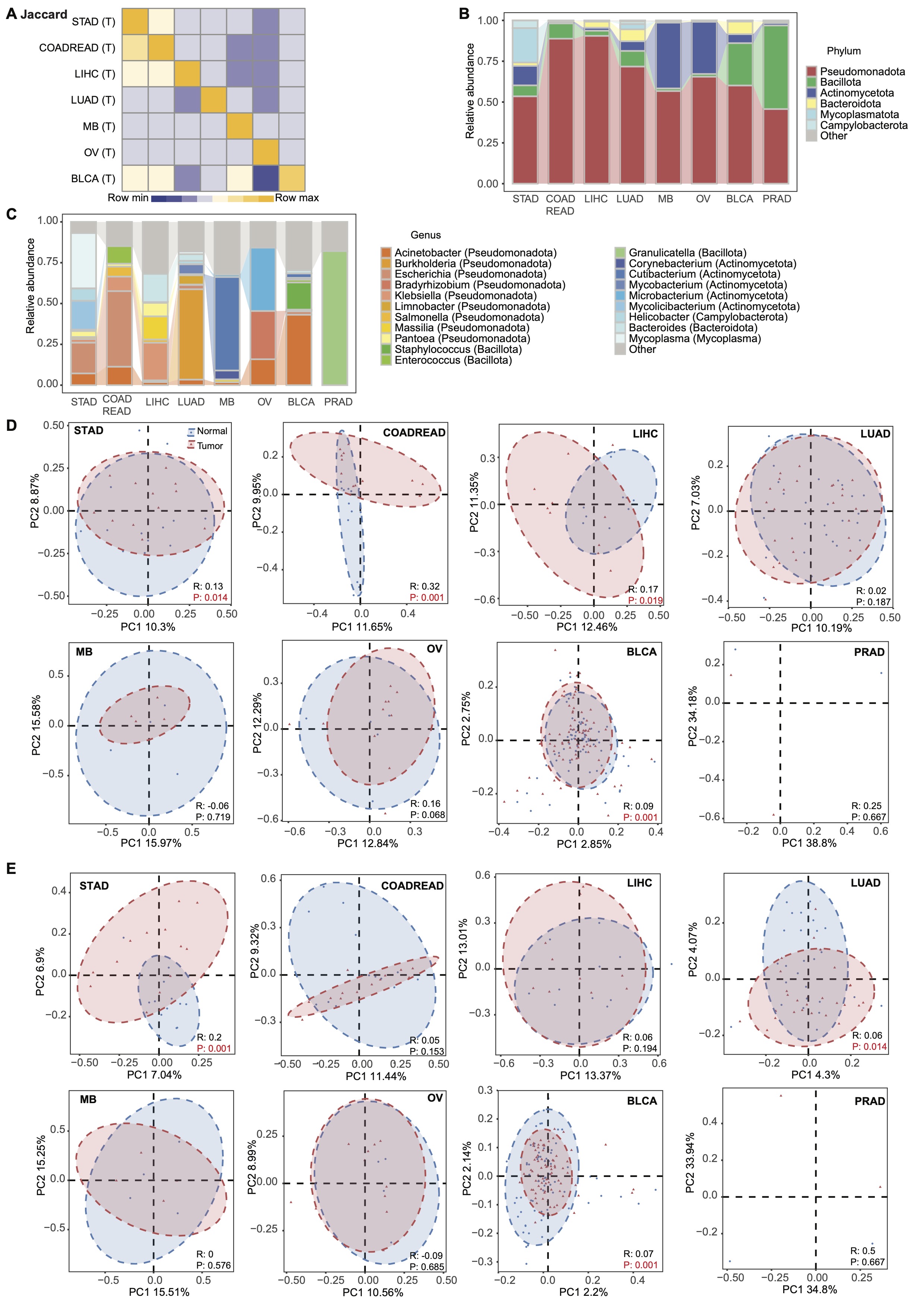

### Supplemental Figure 3

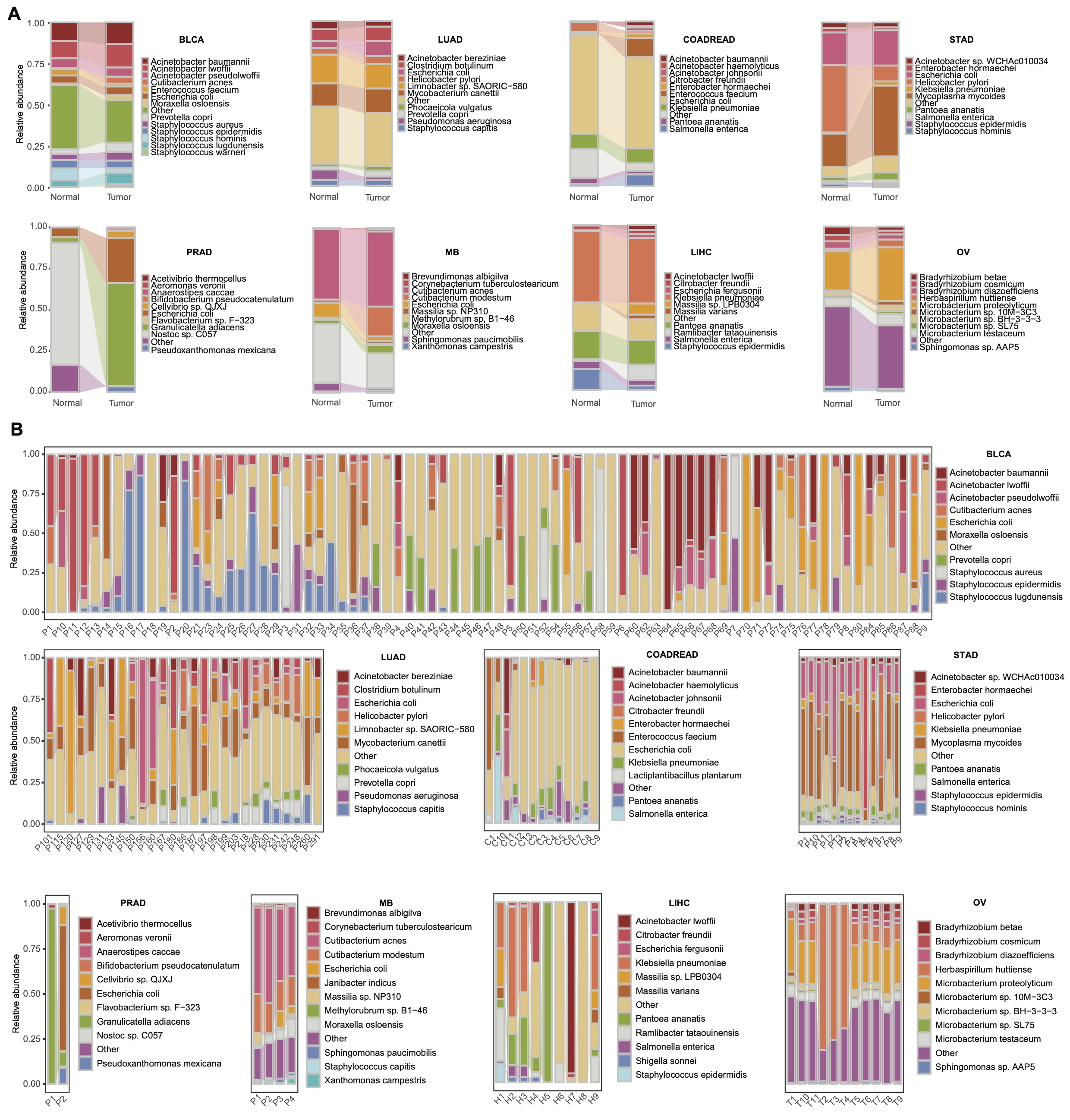

### Supplemental Figure 4

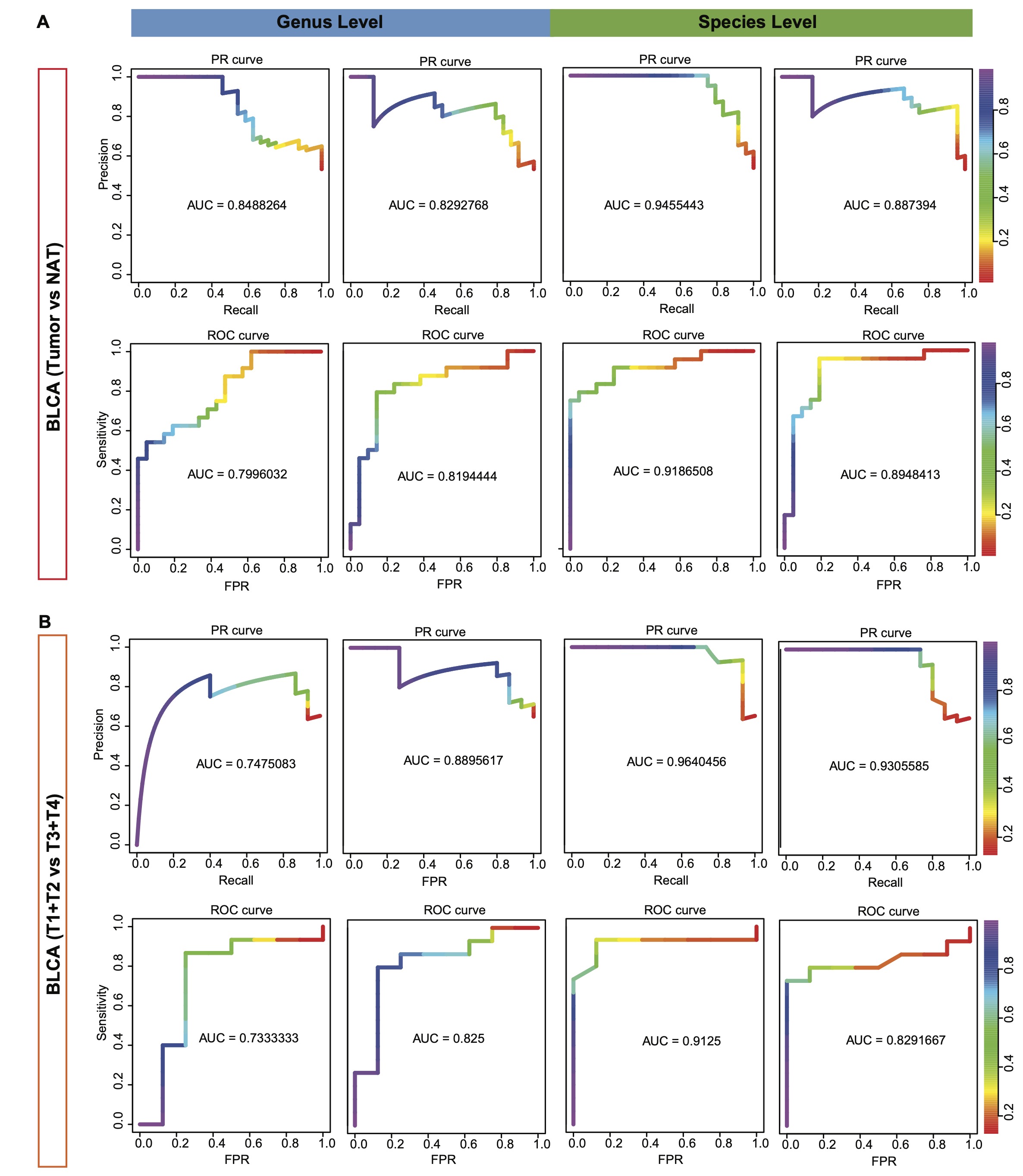

### Supplemental Figure 5

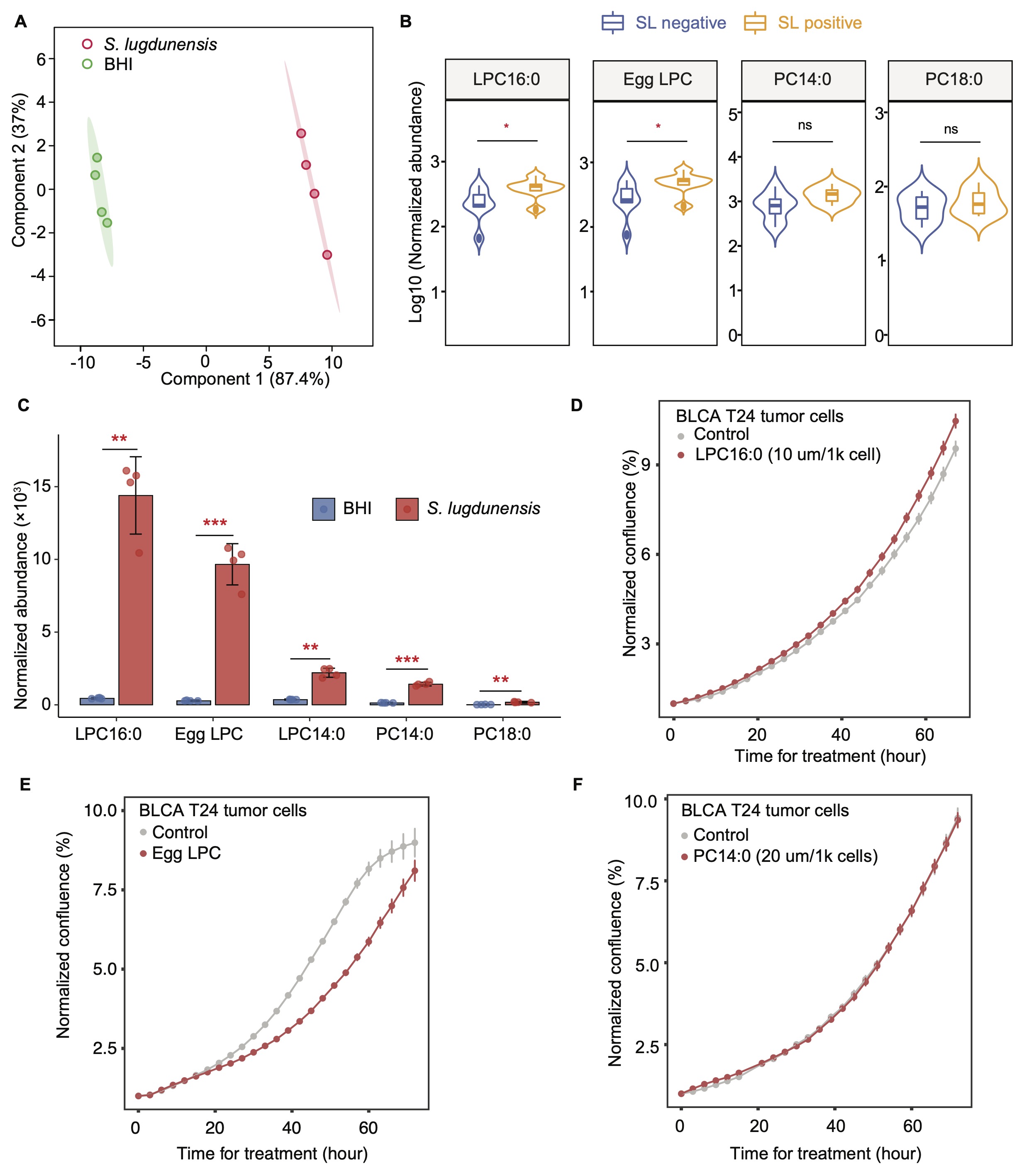
